# mTORC1 activity is essential for disease progression in chronic lymphocytic leukemia

**DOI:** 10.1101/2022.04.19.488778

**Authors:** Natasha Malik, Jodie Hay, Karen M. Dunn, Rinako Nakagawa, Owen J. Sansom, Alison M. Michie

**Author notes:** **Corresponding Author:** Alison M. Michie, Paul O’Gorman Leukaemia Research Centre, Gartnavel General Hospital, 21 Shelley Road, Glasgow, G12 0ZD.

## Abstract

The precise role of mechanistic target of rapamycin complex 1 (mTORC1) during chronic lymphocytic leukemia (CLL) pathogenesis remains to be elucidated. Targeted deletion of mTORC1 component *Raptor* in adult mice reveals that mTORC1 function is essential for initiation and maintenance of CLL. *Raptor*-deficient bone marrow-derived PKCα-KR transduced haemopoietic progenitors failed to generate a CLL-like disease *in vitro*, due to an inability to overcome the mTORC1-mediated block in B cell lineage commitment. Induction of *Raptor*-deficiency in NSG mice transplanted with *Mx1-Raptor* BM-derived PKCα-KR transduced cells after disease was established, revealed a reduced CLL-like disease load and a significant increase in survival in the mice. Interestingly in mice transplanted with an aggressive CLL-like disease, rapamycin treatment reduced disease burden more effectively than AZD2014 (dual mTORC1/2 inhibitor), indicating a skew towards mTORC1 sensitivity with more aggressive leukemic disease. Rapamycin efficiently targeted the translation elongation axis eEF2/eEF2K downstream of mTORC1, resulting in eEF2 inactivation through induction of eEF2^T56^ phosphorylation. Rapamycin treatment of primary CLL cells halted proliferation, modulated eEF2K/eEF2 phosphorylation and inhibited MCL1 expression. Our studies demonstrate that mTORC1 plays an essential role in leukemia progression *in vitro* and *in vivo* in our CLL mouse model, with evidence for increased rapamycin sensitivity in aggressive secondary CLL transplants. Furthermore, the suppression of translation elongation through inactivation of eEF2 may offer a novel therapeutic target for blocking CLL progression.

## Introduction

The clinical management of chronic lymphocytic leukemia (CLL) patients has been transformed by the introduction of targeted therapies that disrupt tumor microenvironmental signals, leading to enhanced survival rates of poor-prognostic patients, highlighting a potential curative strategy (*1*). However, these treatments are not suitable for all CLL patients, and the development of drug resistance has been demonstrated (*2, 3*). Thus, there is an unmet clinical need for novel treatments for high-risk CLL patients. The tumor microenvironment within lymphoid organs of CLL patients promotes interaction of the leukemic clone with the stromal niche, antigen and activated-CD4^+^CD40L^+^ T lymphocytes (*4-7*) playing a pivotal role in enabling survival, proliferation and chemoresistance. Inhibiting the signals orchestrating these events is key to disrupting disease progression (*4*).

Mechanistic target of rapamycin (mTOR) is a serine/threonine kinase that plays an essential role in a multitude of distinct cellular functions including proliferation and survival, facilitated by modulation of protein synthesis and metabolic processes (*8, 9*). mTOR functions in two complexes, mTORC1 and mTORC2, which comprise four shared proteins: mTOR, GβL, DEPTOR and TTI1/TEL2 complex. PRAS40 and RAPTOR (rapamycin TOR sensitive) are unique proteins specific for mTORC1, while RICTOR (rapamycin TOR insensitive), mSIN1, and PROTOR1/2 proteins are specific to mTORC2 (*8*). The critical role played by mTORC1 in positively regulating protein synthesis is highlighted by its regulation of the targets p70 S6 kinase 1 (S6K1) and eukaryotic initiation factor 4E-binding proteins (4EBP1). Phosphorylation of 4EBP1 leads to the release of eIF4E enabling 5’ cap-dependent translation of mRNA transcripts. S6K1 also contributes to activation of 5’ cap-dependent translation through eIF4B activation, promotion of ribosome biogenesis through activation of S6, and inhibition of elongation factor 2 kinase (eEF2K), which relieves inhibition of eEF2 leading to an activation of protein translation elongation (*8, 10*). These processes are highly deregulated in transformed cells, to keep up with the demands of increased cell growth and proliferation.

Studies in mouse models have demonstrated that mTOR signalling plays a critical role during multiple stages of B cell development and maturation (*11*). In particular, analysing the role of mTORC1 revealed significant decreases in B cell generation in mouse models with a conditional knockout (cKO) for *Raptor* in the early stages of development, due to a block in B cell lineage commitment (*11-14*). Therefore, *Raptor* ablation on B cells was assessed using B cell specific models (*Mb1*-Cre), which resulted in a profound block at the preB cell stage abrogating B cell maturation, proliferation, germinal centre (GC) reaction and antibody production (*12*). Additionally, cKO *Raptor* in mature B cells (hCD20-*Tam-*Cre) showed a decrease in GC B cells and nascent antibody secreting plasma cells, and the elimination of GCs resulting in a decline in serum-antibodies (*15*). These studies illustrate the specific importance of mTORC1 at multiple stages of B cell maturation and function. The AKT/mTOR axis plays a key role in leukemogenesis as demonstrated in a mouse model of leukemia induced by *Pten-*loss, in which mTORC1 deletion resulted in a significant increase in survival (*16*). Focusing on CLL, recent studies show that the mTOR pathway is differentially modulated in CLL patients from distinct cohorts with an elevation in 4EBP1 phosphorylation in poorer prognostic CLL patients. This was coupled with an elevation in mTOR signalling in an aggressive CLL murine model (*17*). In agreement with these findings, mice treated with mTORC1/2-targeted inhibitors reduced CLL disease load (*17, 18*). While mTORC1-selective inhibitors, or rapalogs, have shown some promise in targeting CLL through their ability to induce cell cycle arrest by modulating the mTOR/S6K pathway (*19-21*), a deeper understanding of the molecular processes that are deregulated in B cell malignancies downstream of mTOR is required to identify further therapeutic avenues. Here, we assessed the precise role of mTORC1 in the pathogenesis of CLL, analysing mTORC1-inducible KO genetically altered (GA) models, using the *Mx1*-Cre-*Raptor* and *CD19*-Cre-*Raptor* mice together with CLL patient samples and uncover a crucial role of mTORC1-mediated signals in regulating protein translation during CLL progression.

## Results

### *Raptor* (mTORC1)-deficiency negatively impacts on normal B cell development

To demonstrate the robustness of *Raptor* excision in hematopoietic cells in the *Mx1*-Cre model, *Mx1*-Cre^+^*Raptor*^*flfl*^ (*Mx1-Raptor* cKO or *Mx1*-cKO) mice were inoculated with poly(I:C) and assessed 5 weeks post treatment, comparing with inoculated *Mx1*-Cre^-^*Raptor*^*fl/fl*^ (*Mx1-Raptor* control or *Mx1*-control) mice. A significant reduction in *Raptor* gene expression was noted (Fig. 1A), together with a significant decrease in *Ebf1* gene expression, but no change in *Pax5* in the BM and spleen of *Mx1-Raptor* cKO mice (Fig. 1B). These data suggest an early block in B cell development in *Mx1-Raptor* cKO mice compared to controls, in agreement with previous studies (*12-14, 16*). Excision was also validated at the protein level with a significant reduction of RAPTOR expression in the spleen, and thymus in *Mx1-Raptor* cKO mice (Fig. 1C, D). Using both the *Mx1-Raptor* and *CD19-Raptor* (*CD19*-Cre^-^*Raptor*^*fl/fl*^ control; *CD19*-Cre^+^*Raptor*^*wt/fl*^ KO model, we analysed the role of mTORC1 in late B cell development. Similar phenotypic results were noted in the *Mx1-* and *CD19-Raptor* mice showing a significant decrease in the percentage of T2, marginal zone progenitor (MZP), MZ, and follicular 2 (fol2) B cells in the spleen compared to their respective controls (Fig. S1 & S2). This suggests an aberration in late B cells upon *Raptor* ablation, in agreement with previously published studies (*12, 13, 15*).

**Figure 1:**
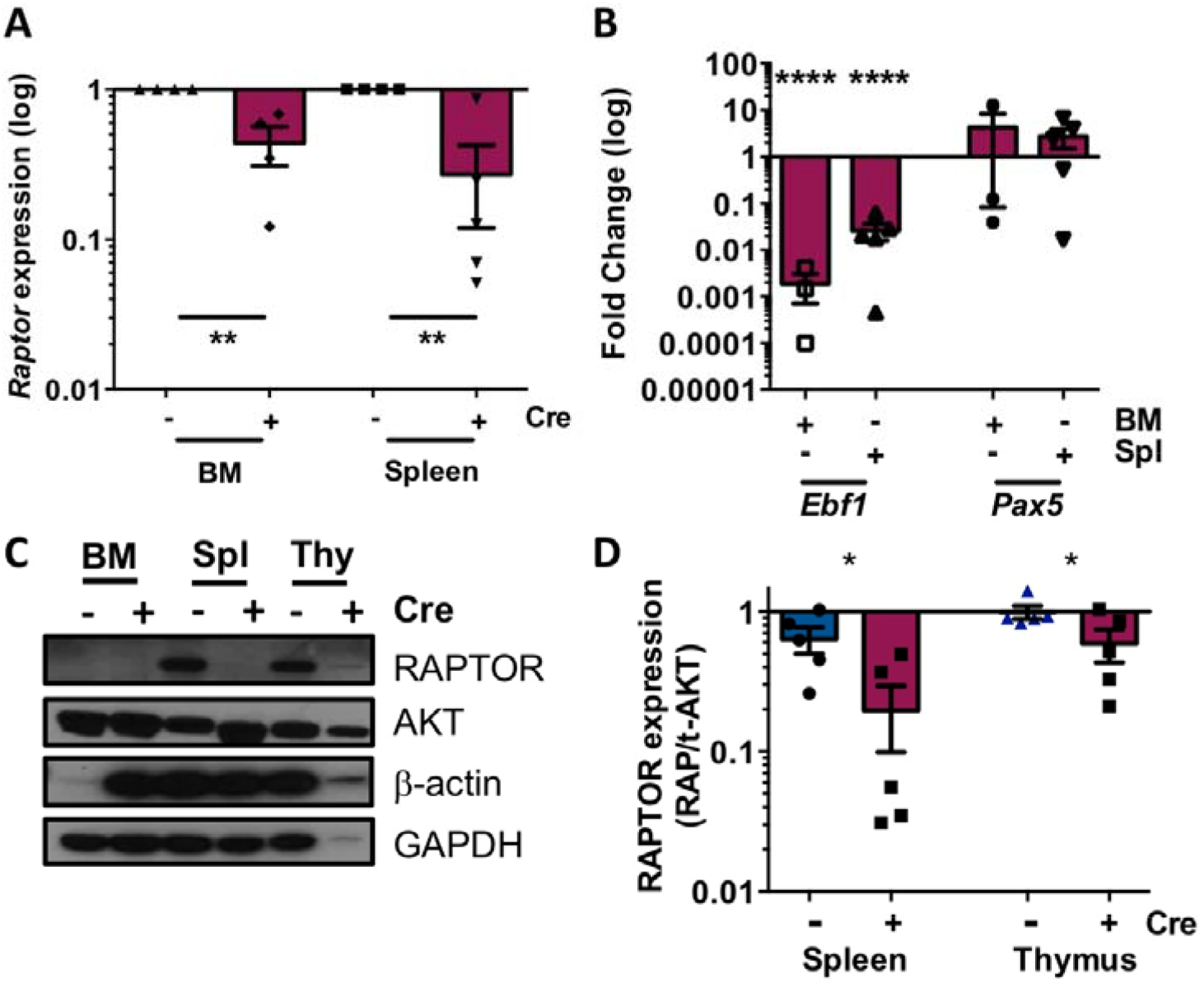
Validation of *Mx1*-cre^+^*Raptor*^*fvfl*^ mouse model for *Raptor*-excision. Gene expression of *Raptor* (n=4) **(A)** and *Ebf1* and *Pax5* (n ≥3) **(B)** in the BM and spleen of *Mx1*-cre^+^*Raptor*^*fvfl*^ mice (red, *Mx1-cre*^+^ *Raptor*^*fvfl*^ cKO. red bar) shown relative to *Mx1-cre Raptor*^*fvfl*^ (*Mx1*-*Raptor* control). Gene expression is shown relative to *GusB* reference gene. **C**. Representative Western blots showing protein expression levels of RAPTOR, with AKT, β-actin and GAPDH as loading controls in *Mx1*-control and *Mx1*-cKO BM, spleen, and thymus extracted from mice. **D**. Quantitative protein expression of RAPTOR in the spleen and thymus of *Mx1*-control and *Mx1*-cKO mice relative to AKT (n≥5 individual mice). All data are expressed as mean ± SEM. p values were determined by two-tailed unpaired *t*-test (p *≤ 0.05; ** ≤ 0.01, p **** ≤ 0.0001).

### mTORC1 plays a fundamental role in CLL-like disease initiation *in vitro*

To assess the role of mTORC1 in CLL initiation, we retrovirally-transduced hematopoietic progenitor cells (HPCs) from the bone marrow (BM) of *Raptor*^fl/fl^ mouse models with kinase dead PKCα (PKCα-KR) to induce a CLL-like disease *in vitro*. Retroviral transduction of *Mx1-Raptor cKO* mice with either MIEV (vector control) or PKCα-KR constructs did not promote the development of B lineage cells or CLL-like disease (GFP^+^CD19^+^ population) respectively, compared to *Raptor*-sufficient control cells (Fig. S3A-C). These findings underlined the essential role played by mTORC1 during B lineage commitment and demonstrates a block in CLL-like disease initiation with *Raptor*-deficiency. Interestingly, *Raptor-* deficiency led to a significant increase in the percentage of CD11b^+^ myeloid cells with time, both in MIEV and PKCα-KR transduced co-cultures, as shown previously in *Raptor*-deficient mouse models *in vivo* ([*12-14*]; Fig. S3A, D, E). In contrast, HPCs derived from *CD19-Raptor* control or KO BM both generated CLL-like disease (GFP^+^CD19^+^ population) after retroviral transduction with PKCα-KR (Fig. S3F, G) suggesting that PKCα-KR is unable to overcome the lineage commitment block induced by *Raptor*-deficiency to induce a CLL-like disease in this model.

### mTORC1 is important for CLL-like disease proliferation *in vitro* and *in vivo*

*CD19-Raptor* KO cells displayed a significant reduction in proliferation when transduced with PKCα-KR compared to *CD19-Raptor* control PKCα-KR cells after 24 hr *in vitro* (Fig. S4A, B). Moreover, there was a significant reduction in migration towards SDF-1 in *CD19-Raptor* KO PKCα-KR cells compared to controls (Fig. S4C, D). These results suggest mTORC1 plays a key role in regulating cellular proliferation and migration in CLL-like cells. To investigate these data further, we assessed the role of *Raptor* excision in CLL-like disease proliferation and maintenance *in vitro* using the inducible cKO model (*Mx1-Raptor*). In the absence of *Raptor* excision, both the *Mx1-Raptor* control and cKO HPCs developed into B cells or a CLL-like disease, and this was maintained similarly over time as expected (Fig. 2A, B). To assess disease maintenance in the absence of mTORC1 *in vitro*, established CLL-like co-cultures were treated with interferon β (IFNβ) to activate the TLR3 receptor and induce the cKO. Confirmation of *Raptor* excision, and concomitant reduction in expression at the gene and protein level in *Mx1-Raptor* cKO PKCα-KR cells compared to *Mx1-Raptor* control PKCα-KR cells, was evident after IFNβ treatment (Fig. 2C-F). Continued co-culture of PKCα-KR cells revealed a significant decrease in the GFP^+^CD19^+^ cell number and reduced proliferation of *Mx1-Raptor* cKO CLL-like cells (and not *Mx1-Raptor* control cells) upon treatment with IFNβ (Fig. 2G-I) indicating an abrogation of CLL-like disease proliferation *in vitro*.

**Figure 2:**
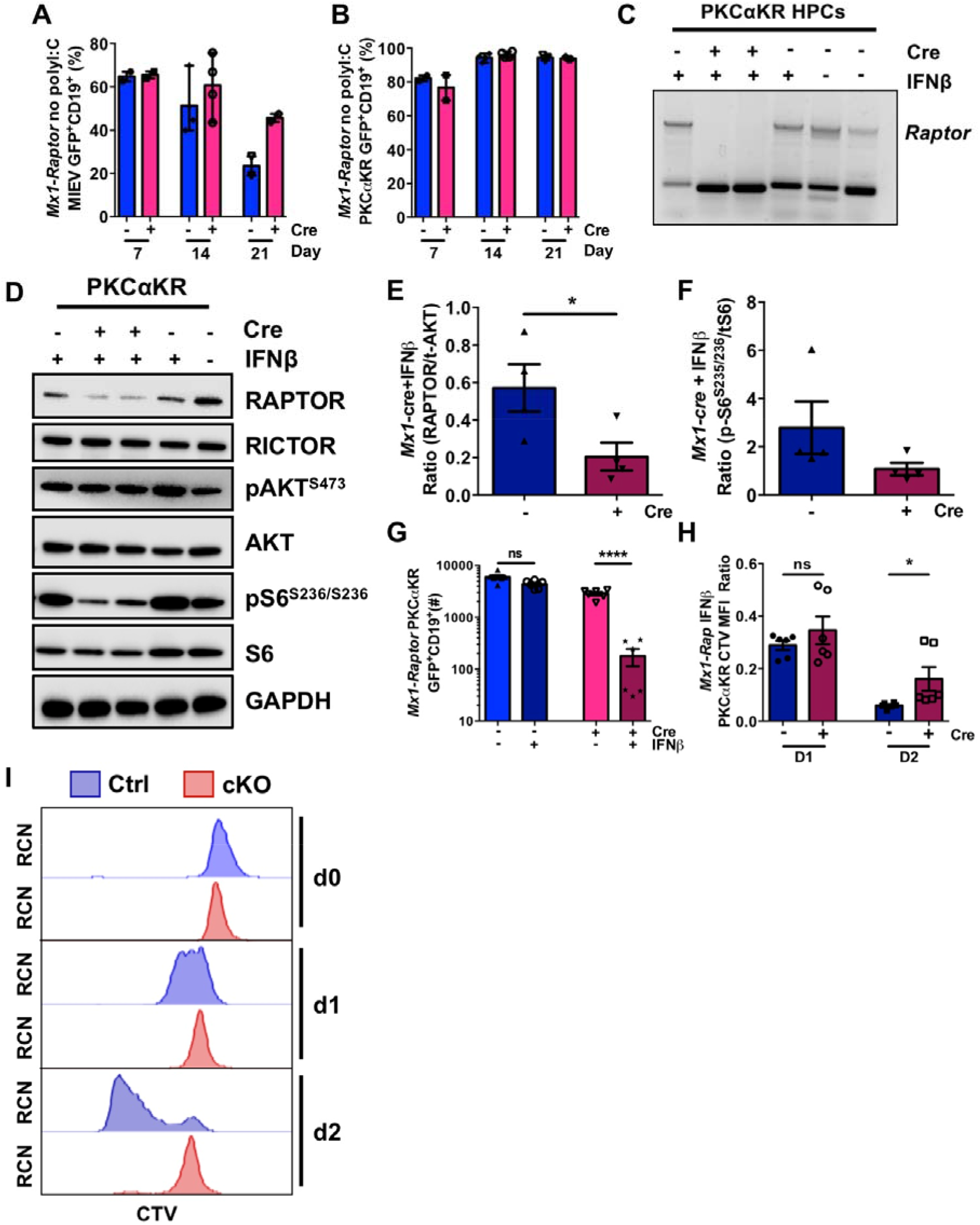
Inducible model of *Raptor*-deficiency abrogates CLL-like disease maintenance *in vitro*. Bar graphs showing the percentage of CD19^+^GFP^+^ *Mx1*-*Raptor* control or *Mx1-Raptor* cKO MIEV **(A)** and *Mx1-Raptor* control or *Mx1-Raptor* cKO PKCαKR **(B)** cells at D7 (n=2), 14 (n=4). and 21 (n=2/3) with no induction of *Raptor* excision. Date expressed as mean ± range of independent cell cultures from individual mice. **C**. DNA agarose gel showing expression of *Raptor* in *Mx1*-*Raptor* control or *Mx1-Raptor* cKO (no polyl:C) PKCαKR cells treated with or without 200U IFNβ for 24 hr and assessed 72 hr post treatment together with *Mx1-Raptor* control HPCs as a reference for *Raptor* expression. Representative Western blot **(D)** and bar graphs showing expression of RAPTOR/AKT **(E)**. pS6^S235/236^/tS6 **(F)** of *Mx1-Raptor* control or cKO PKCαKR (n=4) cells treated with IFNβ and assessed 72 hr post treatment. **G**. Cell number (n-6) of *Mx1*-*Raptor* control or cKO PKCαKR cells either untreated (light blue and pink bars) or treated (blue and red bars) with 200U IFNβ for 24 hr and assessed 72 hr post treatment Bar graphs **(H)** and representative plots **(I)** showing CTV MFI at d0, d1 and d2 of *Mx1*-*Raptor* control or cKO PKCαKR cells (n=6) treated with 200U IFNβ for 24 hr and assessed 72 hr post treatment. Data are expressed as mean ± SEM. p values were determined by two-tailed unpaired f-test (p *≤0.05. p **≤50.001, p***≤0.0001. p ****≤0.00001).

To assess whether mTORC1 plays a similar role in disease maintenance *in vivo*, NOD-SCID-γc^−/−^ (NSG) mice were transplanted with PKCα-KR retrovirally-transduced *Raptor*-sufficient *Mx1-Raptor* control or cKO HPCs. Upon disease establishment (≥10% GFP^+^CD19^+^ cells in blood samples), cohorts of control and cKO mice were treated with poly(I:C) to assess disease maintenance with *Raptor-* deficiency. Weekly blood sampling to screen mice for CLL-like disease demonstrated a reduction in the percentage of GFP^+^CD19^+^ cells in mice transplanted with *Mx1-Raptor* cKO PKCα-KR cells and treated with poly(I:C), while disease load continued to increase in mice transplanted with *Mx1-Raptor* control PKCα-KR cells ± poly(I:C) treatment and *Mx1-Raptor* cKO PKCα-KR cells in the absence of poly(I:C) (Fig. 3A, B & Fig. S5). While no change was noted in splenic weight or total splenic and BM cellularity (Fig. 3C-E), a significant decrease in disease load (GFP^+^CD19^+^ cell number) was seen in the spleens of mice transplanted with *Mx1-Raptor* cKO PKCα-KR cells with poly(I:C) treatment compared to cKO mice without poly(I:C) (Fig. 3F). In addition, there was a significant decrease in the percentage of GFP^+^CD19^+^ cells in the BM and spleen of poly(I:C) treated mice transplanted with *Mx1-Raptor* cKO PKCα-KR cells compared to those transplanted with *Mx1-Raptor* control PKCα-KR cells (Fig. 3G). Comparing the survival of transplanted mice, importantly no significant difference in survival was noted between poly(I:C) treated or untreated mice transplanted with *Mx1-Raptor* control PKCα-KR cells (Fig. 3H), while a significant increase was seen in survival of poly(I:C) treated mice transplanted with *Mx1-Raptor* cKO PKCα-KR cells compared to untreated mice transplanted with *Mx1-Raptor* cKO PKCαKR cells (Fig. 3I). To validate whether these findings were indeed due to *Raptor-*deficiency, protein expression of RAPTOR in the spleen of NSG mice was analysed and a significant reduction was demonstrated (Fig. 3J). These results demonstrate that mTORC1 plays an important role in driving leukemia progression in our CLL mouse model *in vivo*.

**Figure 3:**
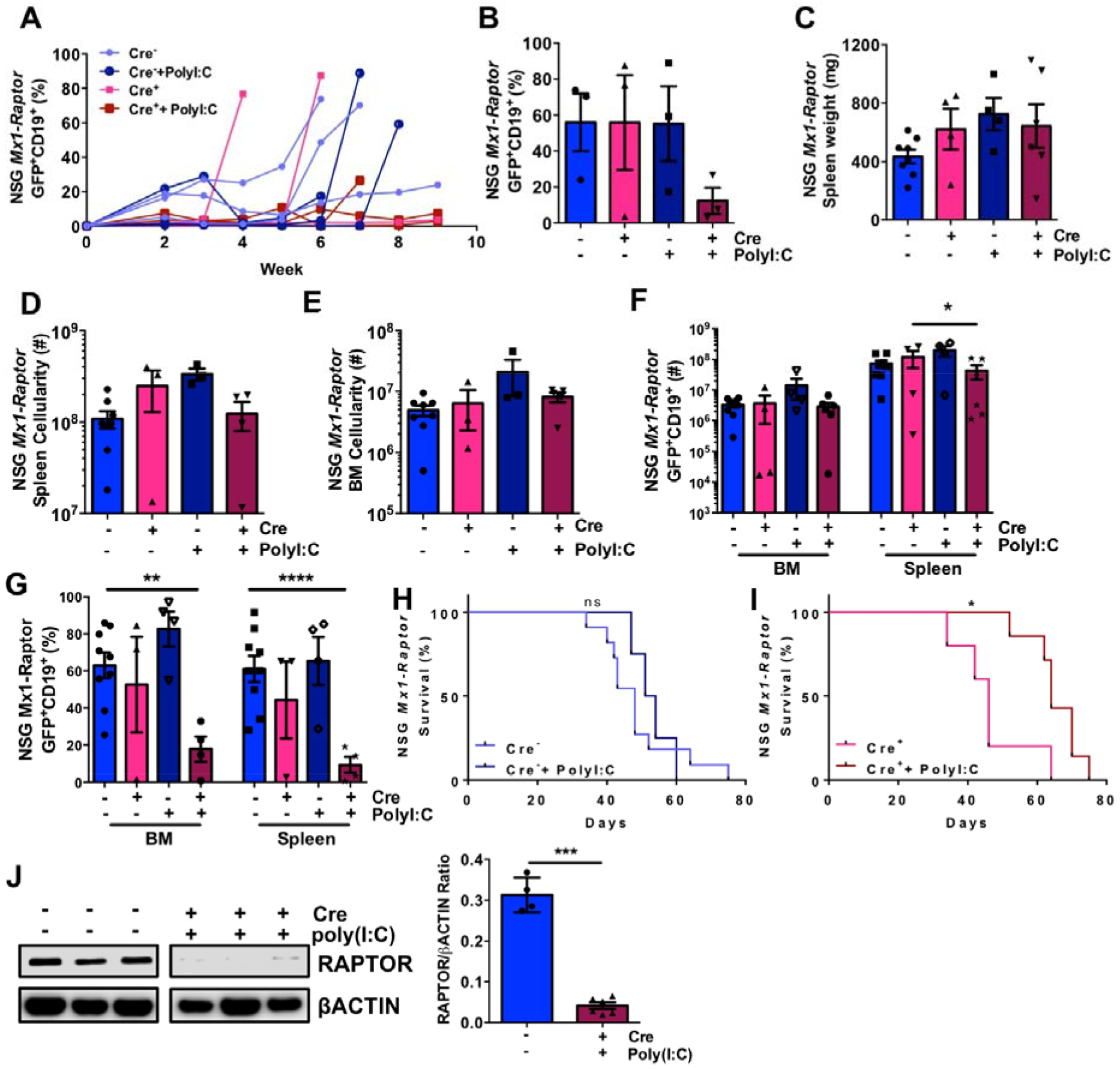
NSG mice with an established CLL-llke disease exhibit a decrease in disease load with Induced *Raptor*-deficiency *in vivo*. NSG mice were Transplanted with *Mx1*-*Raptor*-PKCαKR cells to establish CLL-like disease. Graphs show the percentage of GFP^+^CD19^+^ CLL-like disease in transplanted NSG blood samples taken weekly (representative n=3 Individual mice shown/arm of n>6/arm) **(A)** and at end point **(B)** from *Mx1*-*Raptor* control PKCαKR cells. *Mx1*-*Raptor* control PKCα-KR cells + poly:C after disease development, *Mx1-Raptor* cKO PKCα-KR cells or *Mx1-Raptor* cKO PKCαKR cells + polyl:C after disease development. Spleen weight (mg) **(C)**, spleen **(D)** and BM **(E)** cellularity of NSG mice transplanted with either *Mx1-Raptor* control PKCαKR cells (light blue bar, n=8), *Mx1-Raptor* cKO PKCα-KR cells (pink bar, n=3), *Mx1-Raptor* control PKCαKR cells + polyl:C after disease development (blue bar. n=4), or *Mx1-Raptor* cKO PKCα-KR cells + polyl:C after disease development (red bar, n=5). Cell number **(F)** and percentage **(G)** of GFP^+^CD19^+^ cells in BM and spleen of NSG mice transplanted with either *Mx1*-Raptor control PKCαKR cells (n=8). *Mx1-Raptor* cKO PKCα-KR cells (n=3), *Mx1-Raptor* control PKCα-KR cells + poly I :C after disease development (n=4) or *Mx1-Raptor* cKO PKCαKR cells + polyl:C alter disease development (n=5). Kaplan-Meier survival graphs comparing the percentage survival between NSG mice transplanted with *Mx1-Raptor* control PKCα-KR cells with (n=4) or without (n=11) polyl:C inoculation **(H)**. *Mx1-Raptor* cKO PKCα-KR cells with (n=5) or without (n=7) polyl:C inoculation **(I)** after disease development. **J**. Representative Western blot and densitometry of RAPTOR/pACTIN expression of *Mx1*-*Raptor* control PKCα-KR cells (light blue bar, n=4), or *Mx1*-*Raptor* cKO PKCα-KR cells + polyl:C after disease development (red bar, n=6). These blots are the same as those shown in Figure 7A, therefore showing the same βACTIN loading control. Data are expressed as mean ± SEM. p values were determined by log rank test for Kaplan Meier curves or two-tailed unpaired *t*-test for bar graphs (p *<0.05, p* *<0.001, p * * *<0.0001, p* * * *<0 00001).

To further assess the role of mTORC1 in leukemogenesis, *CD19*-*Raptor* control or KO BM cells were transduced with GFP^+^-PKCα-KR and transplanted into NSG mice. Mice transplanted with *CD19-Raptor* KO cells exhibited a delay in GFP^+^CD19^+^ CLL-like disease burden in the blood compared with *CD19-Raptor* control mice (Fig. 4A), suggesting a non-redundant role of *Raptor* in CD19^+^ cells in disease progression. Although no changes in spleen weight, BM and spleen organ cellularity were observed, there was a significant increase in survival of mice transplanted with *CD19-Raptor* KO PKCα-KR cells compared to controls (Fig. 4B-D). While the *CD19-Raptor KO* PKCα-KR mice survived longer than the controls, disease did eventually accumulate in the mice resulting in similar disease load in the spleen and blood as observed in the control mice. However there was a significant decrease in percentage of GFP^+^CD19^+^ CLL-like cells in the BM, with a similar trend in the LN of mice transplanted with *CD19-Raptor* KO PKCα-KR cells compared to controls (Fig. 4E,F & Fig. S6). This is likely due to an emergence of cells that have escaped *Raptor*-excision, as indicated by the expression of RAPTOR in the spleens of mice transplanted with *CD19-Raptor* KO PKCα-KR cells (Fig. 4G).

**Figure 4:**
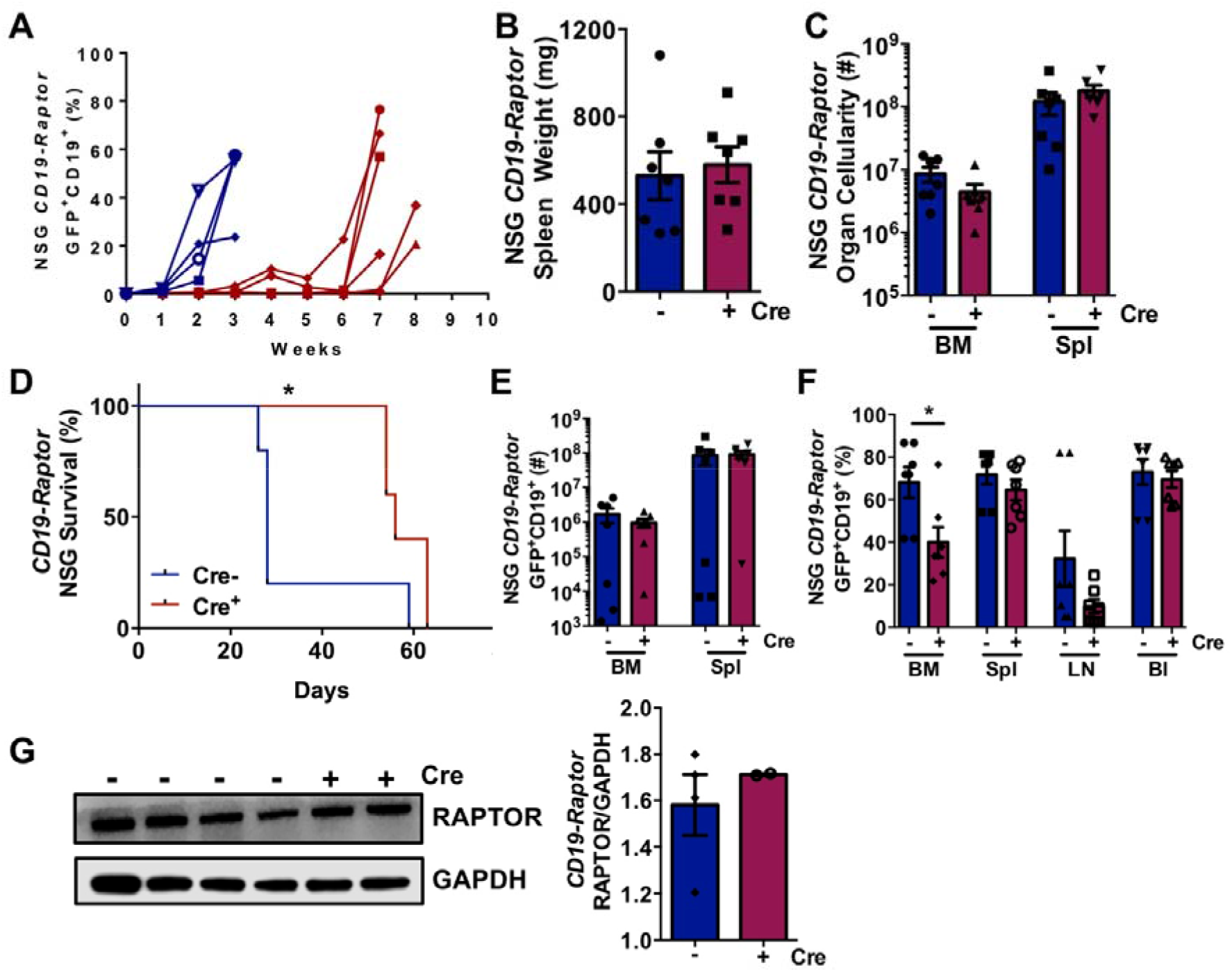
*Raptor*-deficiency in CD19^+^ CLL-like cells increases survival in CLL-like disease *in vivo*. NSG mice were transplanted with PKCα-KR cells and the CLL like disease was established **A**. Weekly blood samples were taken and assessed for percentage of GFP^+^CD19^+^ CLL-like disease in NSG mice transplanted with *CD19*-*Raptor* control (Cre^−^; blue) or cKO (Cre^+^; red) PKCα-KR cells. (n=6 individual mice/arm). Spleen weight (mg) (B) and BM and spleen **(C)** cellulanty of NSG mice transplanted *CD19*-*Raptor* control or cKO PKCα-KR cells (n=7/arm). **D**. Kaplan-meier survival graph comparing the survival of NSG mice transplanted with *CD19*-*Raptor* control (blue) or cKO (red) PKCα-KR cells (n=5/arm). Ccllularity of GFP’CD19^+^ cells in the BM and spleen **(E)** and percentage of GFP^+^CD19^+^ cells in BM, spleen, LN and blood **(F)** of NSG mice transplanted with *CD19*-*Raptor* control {blue) or cKO (red) PKCαKR cells (n=7/arm). Representative western blot **(G)** and protein expression ratio of Raptor and GAPDH (loading control) in the spleen of NSG mice transplanted with *CD19*-*Raptor* control (n=4) or cKO PKCαKR cells (n=2). Data are expressed as mean ± SEM. p values were determined by log rank test for Kaplan Meier curves or two-tailed unpaired *t*-test for bar graphs (p *≤0.05. p **≤0.001).

### Secondary transplantation results in enhanced rapamycin sensitivity of leukemia *in vivo*

To assess the ability of mTOR inhibitors to treat aggressive CLL-like disease, we performed secondary transplants, using splenic cells from mice carrying CLL-like disease (≥95%). Upon disease establishment (≥10% GFP^+^CD19^+^), mice were treated with either vehicle control, rapamycin (4 mg/kg), AZD2014 (15 mg/kg) once daily for 3 wk. Blood samples from mice treated with rapamycin exhibited a clear reduction in GFP^+^CD19^+^ CLL-like cells compared with AZD2014 and vehicle treated mice (Fig. 5A & Fig. S7). A reduction in spleen size was seen in mice treated with rapamycin only, however there were no significant changes in mononuclear cell BM and splenic cellularity (Fig. 5B-D). Focusing on the leukemic cells, a significant reduction in the percentage of GFP^+^CD19^+^ cells was seen in rapamycin treated mice in the BM and blood, with a similar trend in the spleen and LN, with a significant decrease in the percentage of disease burden in the BM, LN and blood of mice treated with rapamycin compared to AZD2014 (Fig. 5E-G). These findings suggest that the CLL-like disease is more sensitive to rapamycin than AZD2014 in secondary transplanted mice.

**Figure 5:**
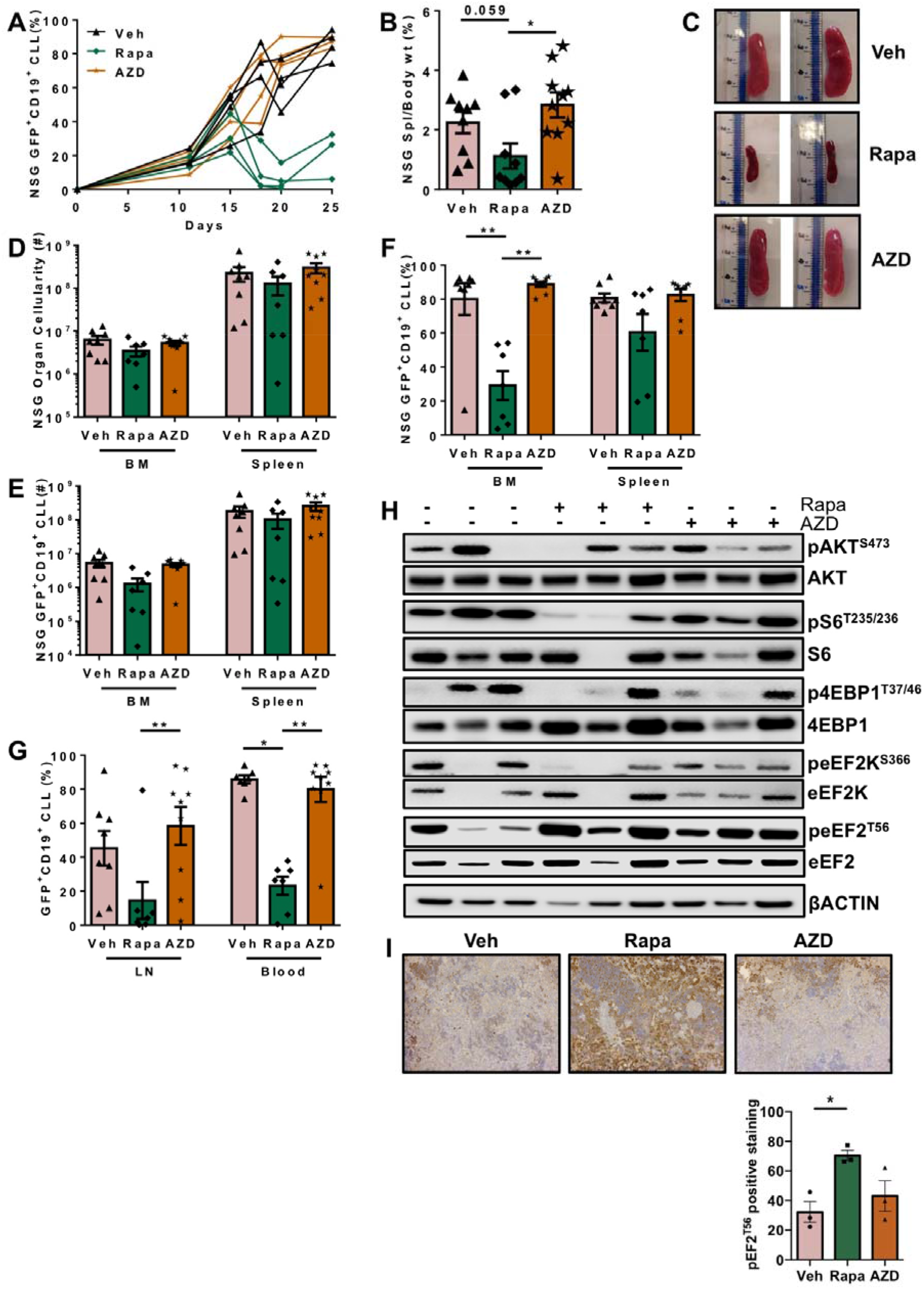
Rapamycin decreases CLL-disease load in an aggressive CLL-like model *in vivo*. Secondary transplants were generated in host mice by transplantation of PKCα-KR cells from spleens of NSG mice with ≥95% CLL-like disease **A**. Weekly blood samples (representative. n=4) were taken and assessed for percentage of GFP^+^CD19^+^ CLL-like disease in mice with established disease and treated with either Veh, Rapa or AZD2014 as indicated. **B**. Percentage of spleen/body weight of drug-treated NSG mice. **C**. Spleens from mice with established disease, treated with vehicle control (Veh; captisol), rapamycin (Rapa) or AZD2014 (AZO) (n≥9). **D**. Total cellularity of the BM and spleen in drug treated NSG mice with a CLL-like disease. Cellularity **(E)** and percentage **(F)** of GFP^+^CD19^+^ CLL-like cells m the BM and spleen together with the percentage **(G)** of GFP’CDW CLL-like cells in the LN and blood of drug treated NSG mice with disease (n≥7). **(H)** Representative western blot of proteins as indicated, using samples of spleens from drug treated NSG mice with disease (n=3). **(I)** Splenic tissue from treated mice were stained for peEF2^T56^ by IHC and the staining was quantified (n=3). Data are expressed as mean ± SEM. p values were determined by a one-way ANOVA (p *≤ 0.05. p ≤ 0. 001).

While there were no significant changes in the phosphorylation status of key mTOR substrates pAKT^S473^, p4EBP1^T37/46^ and pS6^S235/S236^ in spleen from mice treated with drug treated for 3 weeks, we did observe a decrease in pAKT^S473^ in AZD2014 treated mice, and reductions in p4EBP1^T37/46^ and pS6^S235/S236^ signals in both rapamycin and AZD2014 treated mice compared with controls (Fig. 5H, Fig. S8A-C). Analysing further downstream targets of mTORC1 to assess the mechanism driving rapamycin sensitivity, we found that splenic cells from rapamycin-treated mice exhibited a significant decrease in peEF2K^S366^, and an increase in peEF2^T56^ which was significant when analysed by IHC in splenic tissue, compared to vehicle control (Fig. 5H,I & Fig. S8D,E) suggesting possible roles for eEF2 and eEF2K in driving a CLL-like disease in secondary transplants. These data demonstrate that rapamycin more efficiently targets the mTORC1 substrate eEF2K than AZD2014, restoring its ability to inactivate eEF2 and potentially deregulate translation elongation.

### mTORC1-eEF2K/eEF2 signalling is vital for proliferation in primary CLL cells

To assess the role of mTORC1 regulation in human CLL cells, cell cycle status, proliferative capacity and cell viability were investigated in primary CLL patient samples using both the NTL-CD40L/IL21 co-culture method and the MEC-1 cell line in the presence of mTOR targeting inhibitors. In proliferating primary CLL cells, mTOR inhibitors induced cell cycle arrest as indicated by a significant increase in G_0_/G_1_ phase and with a decrease in the S phase in samples treated with rapamycin, AZD8055, a pharmacophore of AZD2014 (Vistusertib) [*22*]), or rapa/Ibr combination as compared to ibrutinib or NDC controls (Fig. 6A-C). Supporting these data, mTOR inhibitors also induced cell cycle arrest in MEC1 cells as indicated by a significant decrease in the S and G_2_/M phases of the cell cycle (Fig. S9A-D). Although a significant reduction in viability was noted upon treatment with rapamycin and AZD8055, mirrored by an increase in early and late apoptosis in MEC-1 cells with AZD8055 treatment (Fig. S9E-H), primary CLL patient samples showed a significant decrease in viability and an increase in apoptosis only in the combination arm (rapa/Ibr) supporting previous studies showing that rapamycin treatment alone does not induce primary CLL cell death ([*17, 20*]; Fig. 6D-F). Finally, we demonstrate a significant decrease in MEC-1 and primary CLL proliferation upon rapamycin, AZD8055, and rapa/Ibr combination treatment (Fig. 6G, H & Fig. S9I, J) compared to NDC. In agreement with the cell cycle data, ibrutinib alone does not inhibit primary CLL cell proliferation, however ibrutinib treatment did reduce MEC-1 proliferation. These studies underscore the importance of mTORC1 in driving CLL proliferation.

**Figure 6:**
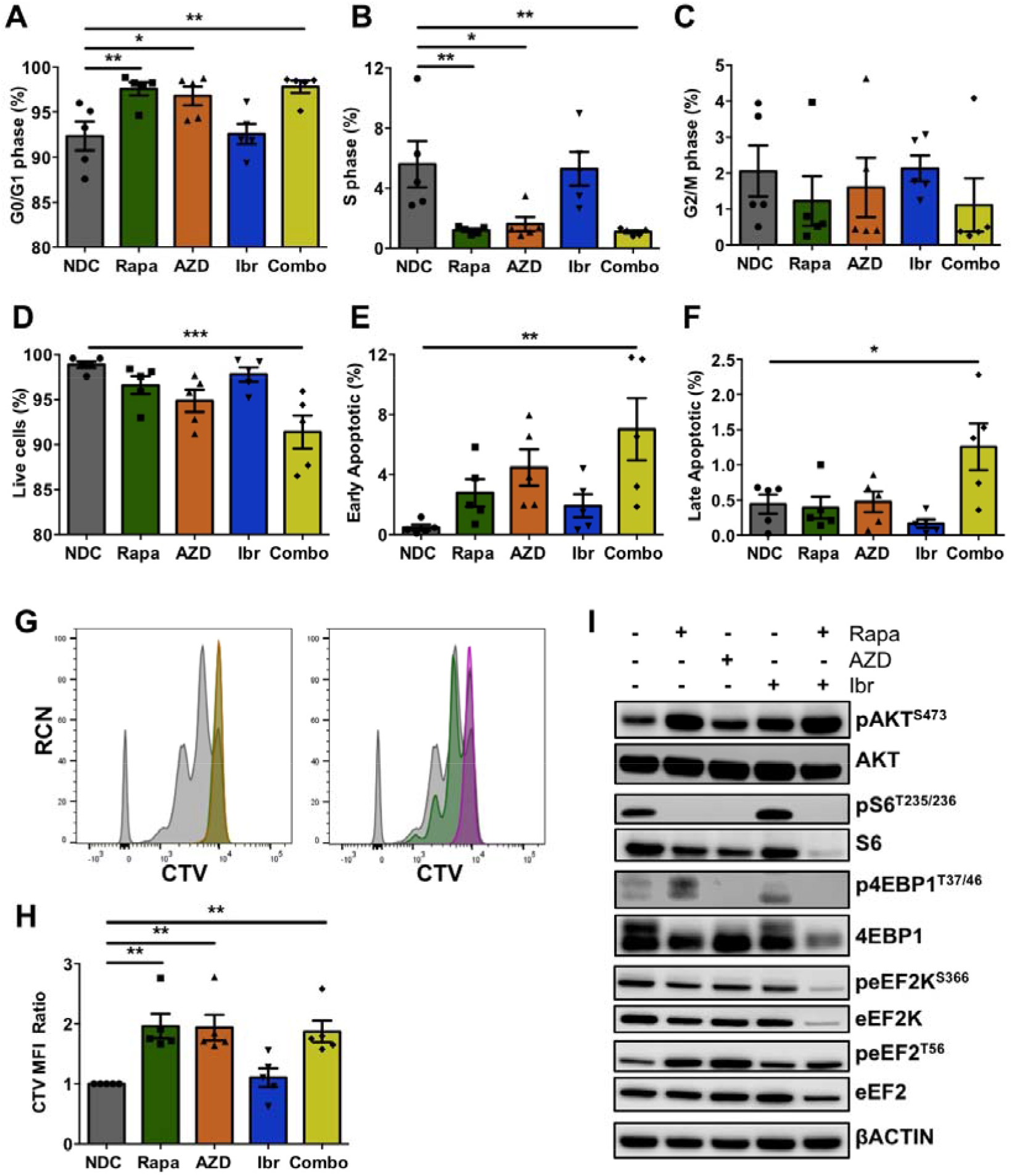
Primary CLL patient cells exhibit altered cell cycling and proliferation with mTOR inhibitors. CLL cells were co-cultured for 5 days with CD40L producing NTL cells with 15 ng/ml IL21 to induce proliferation. Data show averages of 5 individual CLL patients. Graphs showing G0/G1 phase **(A)**, S phase **(B)**. and G2/M phase **(C)** of cell cycling in CLL patient samples treated with either no drug control (NDC, grey), rapamycin (rapa. green). AZD8055 (AZD. orange), ibrutinib (ibr. blue), or a combination of rapamycin and ibrutinib (combo, mustard). Graphs showing a summary of the percentage of live cells **(D)**, early apoptotic **(E)** and apoptotic **(F)** treated primary CLL cells as indicated. Representative flow cytometry plots showing the CTV mean fluorescence intensity (MFI; left = NDC, rapa, AZD; right = NDC, ibr, crombo) **(G)**. and graph showing the CTV MFI ratio **(H)** of CLL patient samples treated as indicated. **(I)** Representative Western blot shown of protein phosphorylation/expression as indicated in primary CLL cells treated with rapamycin (rapa), AZD8055 (AZD). ibrutinib (ibr), or a combination of rapamycin and ibrutinib (combo) or NDC. Blots shown are representative of 5 individual CLL patient samples. Data are expressed as mean ± SEM. p values were determined by a one-way ANOVA (p* ≤ 0.05, p **≤ 0.001).

To investigate the mechanism behind mTORC1-mediated proliferation/survival, we performed Western blotting on lysates generated from proliferating primary CLL cells. As expected, only AZD8055 treatment resulted in significant decreases in pAKT^S473^ and p4EBP1^T37/46^ signals in primary CLL cells, while there was a significant decrease in pS6^S332/336^ with rapamycin, AZD8055, and combo treatment in CLL patient samples (Fig. 6I & Fig. S10A-C). Ibrutinib did not induce a significant impact on mTOR-mediated signals in CLL cells co-cultured with CD40L/IL21 co-culture. Mirroring the phosphorylation events seen in splenocytes of PKCα-KR-transduced *Mx1-Raptor* cKO secondary transplanted mice (Fig. 5H), there was a significant decrease in peEF2K^S366^ in samples treated with mTORC1 inhibitors (rapa and AZD), together with an increase in eEF2^T56^ phosphorylation in CLL patient samples when treated with rapa or AZD, suggesting that targeting mTORC1/S6K may disrupt protein synthesis (Fig. 6I & Fig. S10D, E). To test whether this mechanism was replicated across the disease models assessed, and to analyse the impact of inhibiting the eEF2K/EF2 axis on drivers of proliferation/survival, we assessed the expression of Cyclin A, associated with transition through the S-G_2_/M phase of the cell cycle, and MCL1 associated with increased CLL cell survival, elevation of which is a marker of CLL poor prognosis (*23*). Similar inverse patterns of eEF2K/eEF2 phosphorylation were noted *in vivo*, in mice transplanted with PKCα-KR cells and then treated with polyI:C to delete *Raptor*, or left untreated (Fig. 7A). Further analysis of these mice revealed a decrease in Cyclin A expression. This inverse pattern was also replicated in both PKCα-KR co-cultures and primary CLL cells co-cultured on NTL-CD40L/IL21, which upon treatment with mTORC1 inhibitors inactivated the eEF2K/eEF2 axis and significantly reduced MCL1 and cyclin A expression (Fig. 7B-D), which together with the reduction in pRb^S807/811^ intensity supported mTORC1 signalling playing an integral role in regulating CLL cell proliferation/survival. Collectively these results indicate that translation elongation regulates CLL proliferation and disease pathogenesis.

**Figure 7:**
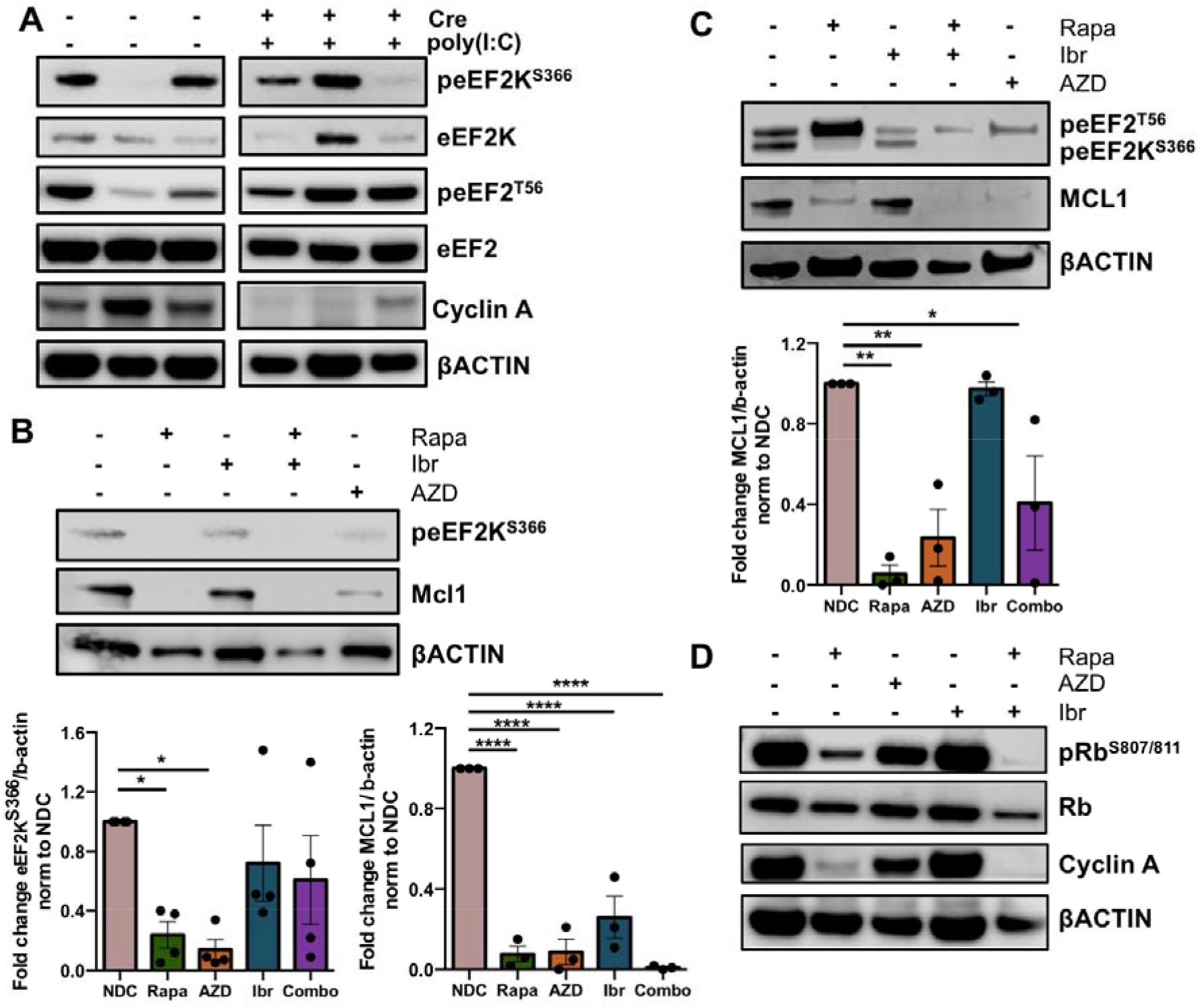
Assessment of mTORC1-regulated protein expression in CLL models. **A**. NSG mice were transplanted with *Mx1*-*Raptor*-PKCαKR cells to establish CLL-like disease. Representative Western blot of indicated proteins in *Mx1*-*Raptor* control PKCαKR cells (n=3), or *Mx1-Raptor* cKO PKCαKR cells + polyl:C after disease development (n=3). These blots are the same as those shown in Figure 3J. therefore showing the same βACTIN loading control. **B**. PKCαKR cells were co-cultured with OP9 cells to establish CLL-like disease *in vitro*. D21 co-cultures were treated with rapamycin (rapa), AZD8055 (AZD), ibrutinib (ibr), or a combination of rapamycin and ibrutinib (combo) or NDC as indicated for 24 hr, and then lysates were prepared Representative Western blot of indicated protein. Densitometry plots of Western blots, with graphs showing peEF2K^S366^/βACTIN (n=4) and MCL1/βACTIN (n=3) relative to NDC. **C&D**. CLL cells were co-cultured for 5 days with NTL-CD40L/IL21 to induce proliferation Representative Western blot showing protein phosphorylation/expression in drug treated primary CLL cells as indicated (n=3). Densitometry plot of Western blots, showing MCL1/βACTIN relative to NDC **(C)**. All data are expressed as mean ± SEM. p values were determined by a one-way ANOVA (p *≤ 0.05. p ** ≤ 0.001, p**** ≤ 0.0001).

## Discussion

We have identified a key role for mTORC1 in CLL maintenance/progression through promotion of proliferation and cell survival, and highlight enhanced sensitivity of aggressive leukemic cells to rapamycin, compared with the dual mTOR kinase inhibitor AZD2014. We highlight for the first time, the importance of the eEF2K/EF2 axis in regulating protein translation elongation downstream of mTORC1-regulated pathways in CLL models, which enable the generation of proteins such as MCL1 and Cyclin A that assist in driving disease pathogenesis. These studies identify a novel therapeutic avenue that may be enable targeting of aggressive leukemia and lymphoma.

We and others have previously established that *Mx1-Raptor* cKO mice displayed significant disruption to B cell development and maturation, at least in part due to a block in B cell lineage commitment at the LSK stage (*11-14*). Furthermore, a similar early block was noted early in T lineage commitment in the absence of mTORC1 (*24*). During the later stages of normal B cell development, significant reductions in T2 B cells were noted, with reductions in MZP, MZ and fol2 populations in the spleen of *Mx1-Raptor* cKO mice, with similar reductions in B cell subsets in *CD19-Raptor* KO mice. These results agree with previous literature determining the function of late B cell populations (*12, 15*). T1 cells transition into T2 cells leading to subsequent fol2 B cells. Fol2 B cells are more primitive, quiescent and less abundant compared to fol2 cells, which form the majority of the recirculating pool. MZP B cells arise from fol2 cells leading to MZ B cells. Fol2 cells also generate fol1 cells (*25*). Therefore, observing a significant decrease in T2 and the primitive fol2 cells and a concomitant decrease in MZP and MZ cells in *Mx1-Raptor* mice is expected due to the fundamental role of mTORC1 in B cell development (*11*).

Combining these *Raptor*-deficient models with our PKCα-KR CLL-like model enabled us to analyse leukemia initiation. PKCα-KR expression was unable to rescue the lineage commitment block caused by *Raptor-*deficiency and therefore failed to initiate the development of CLL in *Mx1-Raptor* cKO-PKCα-KR cells. In an attempt to circumvent this issue, we used the *CD19-Raptor* KO model which promotes *Raptor-*deficiency in CD19^+^ cells, at the preB cell stage after lineage commitment has occurred (*26*). While the cells were able to survive to a similar degree as the *CD19-Raptor* sufficient PKCα-KR cells, *CD19-Raptor* KO PKCα-KR cells exhibited reduced levels of proliferation and migration *in vitro*.

Moreover, IFNβ treatment of *Mx1-Raptor* cKO-PKCα-KR-transduced BM cells to excise *Raptor*, reduced cell proliferation *in vitro*. mTORC1 has also been found to play an essential role in the initiation of T-ALL and AML initiation (*13, 24*), indicating that *Raptor* plays a critical role in initiating haemopoietic malignancies, at least in part by reducing proliferative capacity. In contrast, disease maintenance was significantly negatively impacted upon excision of *Raptor in vivo* after leukemia had been established, which was coupled with an increase in survival in our CLL models. Of interest, Kalaitzidis *et al*., demonstrated that *PTEN*-loss leads to MPN due to mTORC1 activation, which can be reversed by inducing *Raptor* loss after MPN, suggesting a fundamental role of mTORC1 in leukemia progression (*16*). *CD19-Raptor* KO PKCα-KR transplanted mice also displayed increased in survival in mice compared to *CD19-Raptor* control PKCα-KR transplanted mice. However, this increase in survival was not sustained, with the mice dying at a later timepoint with an increase in disease load in the blood of NSG mice with *CD19-Raptor* KO PKCα-KR cells. Further assessment revealed a re-emergence of RAPTOR expression suggesting that while *Raptor* excision does abrogate disease progression, the *CD19*-Cre-loxP model does not induce an efficient knockout in CLL-like cells leading to ‘escaped deletions’ and subsequent re-population of disease. Indeed, the *CD19*-Cre model has previously been demonstrated to induce excision with 80-93% efficiency (*27*).

We have previously shown that transplantation of PKCα-KR CLL-like cells into mice leads to a CLL-like disease *in vivo* and targeting the disease with either rapamycin, AZD8055, AZD2014 or ibrutinib was not as potent as a combination therapy of AZD2014 and ibrutinib (*17*). Indeed, AZD8055 more robustly and significantly reduced disease compared with rapamycin in primary PKCα-KR transplanted mice (*17*). Using a more aggressive leukemia model by performing secondary transplants revealed that AZD2014 was inferior compared to rapamycin in decreasing disease load *in vivo*. This was evident with tumor load in the blood during treatment, splenic size and proportion of leukemic GFP^+^CD19^+^ cells in the organs. Collectively, these studies indicate that the secondary transplanted disease is more rapamycin/mTORC1-sensitive. Similar findings have been noted in an APC-deficient mouse model of colorectal cancer (CRC), in which mTORC1 targeted inhibition via rapamycin treatment reduced tumor growth and increase survival of mice (*28*). Of note, a phase II clinical trial showed similar results in patients with refractory renal cancer, with AZD2014 treated patients exhibiting a shorter progression-free survival and increased disease progression compared to patients treated with everolimus, a compound related to rapamycin (*29*). Therefore, delineating the distinct mechanism of action of the two drugs may enable better treatment options for cancer patients.

Elevated protein synthesis is essential for driving oncogenesis, enabling increased cellular proliferation and survival of cancer cells through the generation of proteins such as MCL1 and MYC, and proteins responsible for regulating the cell cycle. mTORC1 substrates play a central role in regulating protein synthesis, with studies demonstrating that suppression of translation limits tumorigenesis (*10, 28, 30, 31*). Through phosphorylation of 4EBP1, mTORC1 inhibits its interaction with eIF4E thus enabling the generation of the eIF4F complex that mediates cap-dependent translation. We have previously shown that *ex vivo* peripheral blood-derived poor prognostic CLL cells have upregulated p4EBP1, suggesting that these cells possess elevated cap-dependent translation (*17*). This is supported by studies demonstrating that B cell receptor crosslinking on CLL cells resulted in elevated expression of eIF4A and eIF4GI (part of the eIF4F complex) and drives MYC and MCL1 translation (*32, 33*). Further regulation of protein translation is mediated by mTORC1/S6K activation, through phosphorylation and inhibition of eEF2K which in turn negatively regulates eEF2 function (*31, 34*). eEF2 promotes translation elongation, however eEF2K-mediated phosphorylation at residue T56 results in excluding EF2 from the ribosome, thus suppressing translation elongation (*35, 36*). Our CLL models demonstrated that the eEF2K/eEF2 signalling axis was inhibited by mTOR inhibitors and that our aggressive secondary transplant leukemia model was particularly rapamycin sensitive, identifying the possibility that poor prognostic leukemias may benefit from therapeutic agents that target translation elongation. Indeed, analysis of the splenic tissue from secondary transplants demonstrated a significant downregulation in eEF2K^S366^ and upregulation of peEF2^T56^, similar to that noted in CRC patient tumor samples, where low eEF2K activity has been associated with a worse prognosis of patients (*31, 37*). These findings were mirrored in proliferating primary CLL samples treated with rapamycin. Importantly, the inverse regulation of eEF2K/eEF2 phosphorylation was also evident in proliferating primary CLL samples, and we demonstrated that induction of peEF2^T56^ through mTORC1 inhibition resulted in reduced cyclin A and MCL1 expression.

Collectively, we demonstrate that mTORC1 plays a fundamental role in CLL initiation and progression *in vitro* and *in vivo*. These data identify a novel mTORC1-biased mechanism by which the PKCα-KR murine CLL-like disease is regulated and suggests that selective inhibition of protein synthesis and elongation processes may represent a promising therapeutic avenue in patients with aggressive CLL These studies highlight the potential for eEF2K^S366^/eEF2^T56^ as novel biomarkers for screening leukemia/lymphoma patients to identify those that would be responsive to rapalogs.

## Materials and Methods

### Mice, primary cells and cell lines

Mice expressing the Cre/loxP system with *Raptor* (*Raptor*^*fl/fl*^) were obtained from Prof. Michael N. Hall (University of Basel, Switzerland)(*38*) and crossed with *Mx1*-Cre^+/-^ or *CD19*-Cre^+/-^ (from Dr. Dinis Calado (Francis Crick Institute, London, UK)) GA mice, to obtain the desired KO models. B6.SJL mice (6-10 wk old) were used as a source of wild type (WT) BM and NSG immunocompromised mice were used as hosts (6-8 wk old) for transplantation of CLL-like disease. All mice were housed at the Beatson Research Unit (BRU; Glasgow, UK) or the Veterinary Research Facility (VRF, Glasgow, UK). All experimental protocols were approved by the local AWERB committee and national Home Office (PD6C67A47), and all methods were carried out in accordance with standard animal housing conditions under local and UK Home Office regulations. The *Mx1* promoter is activated upon TLR3 activation (*26*) by inoculating cre^+^-*Raptor*^*fl/fl*^ mice with 10 mg/kg TLR3 agonist poly(I:C) once every two days as indicated (GE Healthcare, WI) to induce *Raptor* cKO in the mouse. *CD19* is expressed at the proB cell stage of B lineage development, thus targeting *Raptor-*deficiency in the B lineage only (*39*). Blood, spleen and lymph node (LN) were removed from transplanted mice and organs were crushed to obtain a single cell suspension. Cells collected from the BM and spleen were enriched for haemopoietic cells by density centrifugation using Lympholyte-Mammal (Cedarlane, Canada) as described previously (*14*). Peripheral blood was obtained, after informed consent, from patients with a confirmed diagnosis of CLL that were treatment-naïve or had received treatment but not in the preceding 3 months. The studies were approved by the West of Scotland Research Ethics Service, NHS Greater Glasgow and Clyde (UK) and all work was carried out in accordance with the approved guidelines (REC Ref: 20/WS/0066). Linked clinical data of prognostic markers of the CLL cohorts were recorded (Table S1). CLL cell purity was ≥ 95% in all cases, determined by flow cytometry. Primary CLL cells and the MEC-1 CLL cell line were cultured at 1×10^6^/mL in RPMI-1640 containing 10% foetal bovine serum (FBS), 50 U/mL penicillin, 50 mg/mL streptomycin, and 2 mM L-glutamine (CLL medium; Invitrogen Ltd., Paisley, UK). OP9 stromal cells were maintained in OP9 medium (αMEM medium supplemented with 20% FBS, 50 U/mL penicillin, 50 mg/mL streptomycin, 2 mM L-glutamine, 10 mM HEPES, 1 mM sodium pyruvate, 10 µg/ml gentamycin, 50 µM βME). The fibroblast cell line constitutively expressing CD40L (NT-L-CD40L) and the non-transduced cell line (NT-L) were maintained in CLL medium.

### Retroviral Transduction of haemopoietic progenitors, transplantations and drug treatments

BM-derived HPCs isolated from *Mx1*-Cre *Raptor* (± poly(I:C) inoculation), *CD19*-Cre *Raptor* mice or B6.SJL WT mice, were processed and re-suspended in 80 µl MACS buffer (2% FBS, 2 mM EDTA in PBS) and 20 µl CD117 MicroBeads (MACS Miltenyi Biotec, Surrey, UK) to enrich LSK cells as per the manufacturers protocol. Thereafter the LSKs were retrovirally-transduced with either empty vector control (MIEV) or kinase dead PKCα (PKCα-KR) vector to induce a CLL-like disease as described previously (*40*). Cells were co-cultured with the stromal OP9 cell line supplemented with IL-7 and Flt-3 until day 7-10 prior to transplantation, passaging every 2–3 days. The cells were cultured at 37ºC in a humidified incubator with 5% (v/v) CO_2_. 5×10^5^ cells/100 µl cells were transplanted into NSG host mice via tail vein injections to establish a CLL-like disease *in vivo*. Disease progression was monitored by blood sampling. Once disease was established in host mice transplanted with *Mx1-Raptor* control/cKO PKCα-KR cells, half the cohort was given 4 doses 10 mg/kg poly(I:C) inoculations over 8 days to induce *Raptor*-deficiency. Secondary transplants were performed using splenic cells isolated from mice transplanted with PKCα-KR-transduced cells. A single cell suspension of spleens was generated under sterile conditions and 3×10^5^ splenic cells were transplanted into NSG host mice as secondary transplants. After confirmation of CLL-like disease (≥10% GFP^+^CD19^+^ cells in the blood), host mice were treated for up to 3 wk with mTOR inhibitor AZD2014 (a gift from AstraZeneca, Cambridge, UK), rapamycin or vehicle control. The drugs were formulated as follows: AZD2014 - 3 mg/mL in 20% Captisol (Ligand Pharmaceuticals, Inc., La Jolla, CA) and administered daily at 15 mg/kg via oral gavage (OG); Rapamycin was dissolved in Tween-80 5.2%/PEG-400 5.2% (v/v) and delivered once daily by intraperitoneal (ip) injection at 4 mg/kg (*17*).

### In vitro cKO induction

Retrovirally-transduced HPCs derived from *Mx1*-*Raptor* mice were co-cultured with OP9 cells *in vitro* until d10. Then 0.5-1×10^6^ cells/well were treated with 200 U/well IFNβ for 24 hr to induce excise *Raptor*. The cells were harvested 4 days post treatment and used in ongoing experiments, as indicated.

### Primary CLL proliferation co-cultures

CLL cells were co-cultured with the NTL-CD40L cell line at a ratio of 1:75, as described previously (*41, 42*). The culture was supplemented with 15 ng/mL IL21 (PeproTech, UK) to aid the induction of proliferation. These co-cultures were treated with DMSO as vehicle/no drug control (NDC), 10 nM rapamycin, 100 nM AZD8055, 1 µM ibrutinib or a combination of rapamycin and ibrutinib, as indicated.

### Flow Cytometry

Single cell suspensions from *in vivo* or *in vitro* experiments were prepared for phenotypic analysis by flow cytometry as described previously (*17*). All antibodies were purchased from BD Biosciences (Oxford, UK), except CD21 (Clone: 7E9, BioLegend), CD1d (Clone: 1B1, BioLegend), IgD (Clone: 11-26c.2a, BioLegend), CD23 (Clone: B3B4, BioLegend). For cell cycle analysis, treated 1×10^6^ PKCα-KR and 2×10^6^ primary CLL patient cells were fixed and permeabilized by adding 1 ml cold 80% ethanol dropwise to the pellet and stored at -20^°^C until analysis. After washing the samples, 350-500µl propidium iodide (PI)/RNAse Staining Buffer (BD Biosciences) was added to each sample and incubated at RT in the dark for 15 min before analysis. For cell proliferation, cells were stained with CellTrace™ Violet stock solution (Invitrogen, Paisley, UK) as per the manufacturer’s protocol. PKCα-KR cells were acquired for 3 consecutive days (every 24 hr) and primary CLL patient samples were recorded on day 5 of culture to assess proliferation. For apoptosis analysis, treated 5×10^5^ PKCαKR or primary CLL patient cells were washed with 1x HBSS (ThermoFisher Scientific) at 300*g* for 5 min at RT. Cells were incubated in 100 µl HBSS containing 2.5 µl Annexin V and 2.5 µl 7AAD (BD Biosciences) for 10 min at RT in the dark and analysed. All data were acquired on a FACSCantoII with BD FACSDiva software and analysed using FlowJo software (Tree Star Inc., OR).

### qRT-PCR

RNA was extracted from fresh cells following the RNAeasy Qiagen Kit protocol. cDNA was made using standard protocols (Invitrogen). qPCR was carried out on a 384 well plate in triplicate by the 7900HT Fast Real-Time PCR system (Applied Biosystems, Warrington, UK). The primers used are listed in Table S2.

### Western blotting

Protein lysates of cell pellets were prepared in lysis buffer (20 mM Tris pH 7.4, 2mM EDTA, 1% Triton, 1mM DTT) containing protease inhibitor cocktail and phosphatase inhibitor cocktail (Roche) and incubated on ice for 30 min. Bradford assay was performed to quantify protein samples using a standard protocol (Bio-Rad). 4-12% pre-cast gels (NuPAGE Novex BisTris) were used for gel electrophoresis. Western blotting was performed by following the standard protocol provided (Invitrogen). The PVDF membranes were blocked in 5% milk and primary antibodies used were prepared in 1% BSA. List of antibodies are listed in Table S3. Signals were detected using Imobilon Forte HRP substrate (Millipore).

### Statistics

Statistical analyses were carried out using GraphPad Prism 6 Software (GraphPad Software Inc., San Diego, CA). p values were determined by two-tailed students paired or unpaired *t*-test, log rank test for Kaplan Meier curves or mixed model ANOVA on a minimum of at least 3 biological replicates as indicated. Biological replicates were derived from individual mice or cell culture conditions from distinct biological samples. Mean *±* standard error of mean (SEM) is shown. * p≤0.05; ** p≤0.01; *** p≤0.001; **** p≤0.0001.

## Acknowledgements

The authors thank the CLL patients who kindly donated blood samples to this study. We are also grateful to Catherine Winchester (BICR) for critical appraisal of the manuscript.

## Author Contributions

Conceptualization: AMM, OJS, NM, JH

Methodology: NM, JH, KMD, AMM, RN

Investigation: NM, JH, KMD, AMM

Funding acquisition: AMM, OJS

Project administration: AMM

Supervision: AMM, OJS

Writing – original draft: AMM, NM

Writing – review & editing: NM, JH, KMD, RN, OJS, AMM

## Data and materials availability

All data are available in the main text of the supplementary materials.

