## Supplementary material for "mTORC1 activity is essential for disease progression in chronic lymphocytic leukemia": Suppl Figures 1-10 * Suppl Tables 1-3

Supplementary Method:

*Migration Assay*: Cells (2-5x10^6^ cells/ml) were cultured for 2 hr at 37ºC in resting medium (DMEM containing 0.5% bovine serum albumin (BSA), 10 mM HEPES, 1 mM sodium pyruvate, 100 µg/ml streptomycin, 100 U/ml penicillin, 2 mM L-glutamine). Resting medium was supplemented with 150 ng/ml SDF-1 (PeproTech, UK) to make migration medium. Transwell^®^ permeable support chambers (Corning Inc, ME, USA) were set up such that the bottom contained 600 µl migration medium and 100 µl cells from the resting step was pipetted onto the top chamber. Each condition was carried out in technical duplicates. For negative and positive controls, resting medium was used instead of the migration medium: rested cells were either pipetted onto the chamber or directly into the bottom respectively. Cells were cultured at 37ºC for 4 hr. 150 µl medium was pipetted from every well in technical triplicate and counted in the flow cytometer on low for 30 sec. Data shown is an average of 3 individual mice.

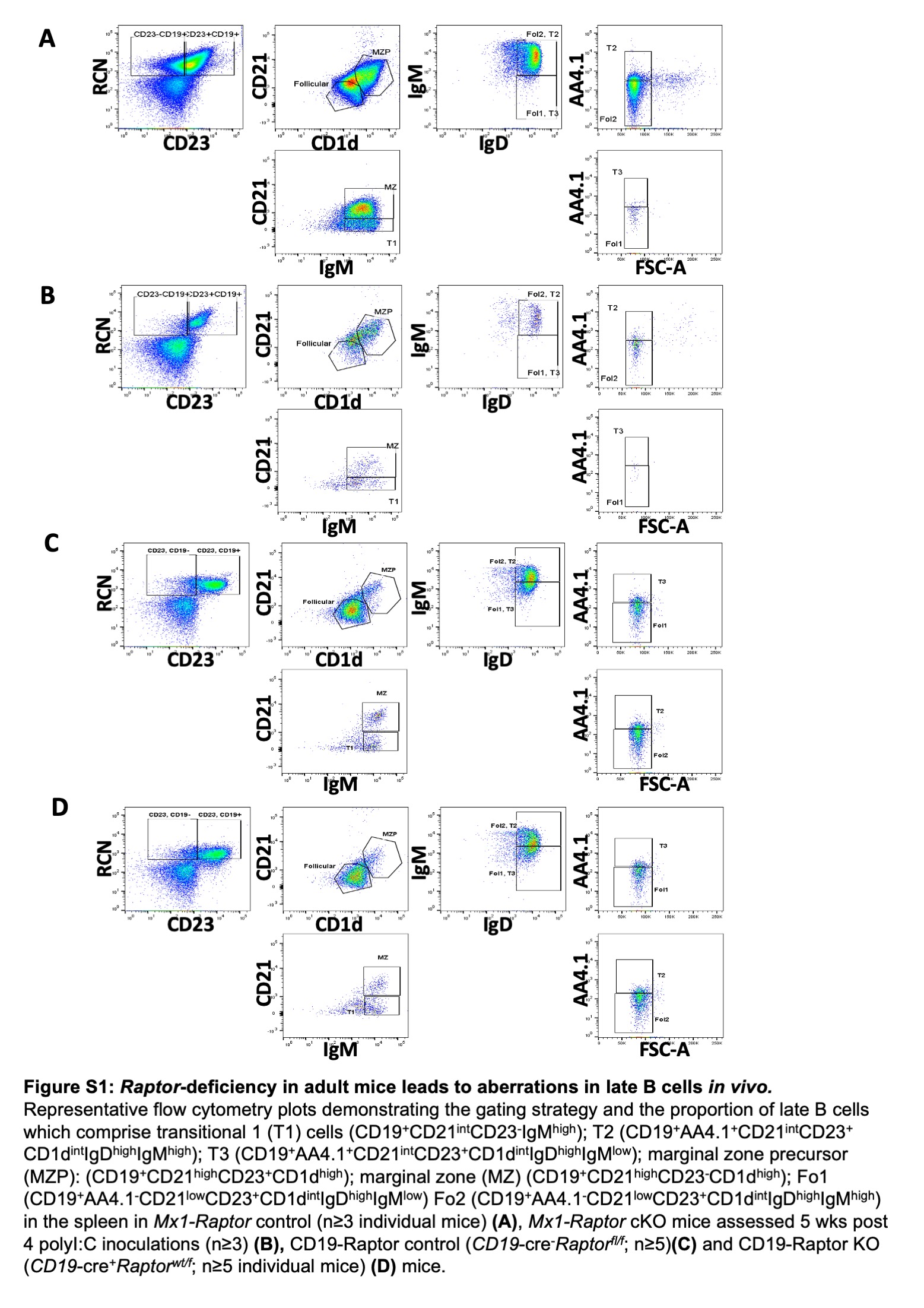

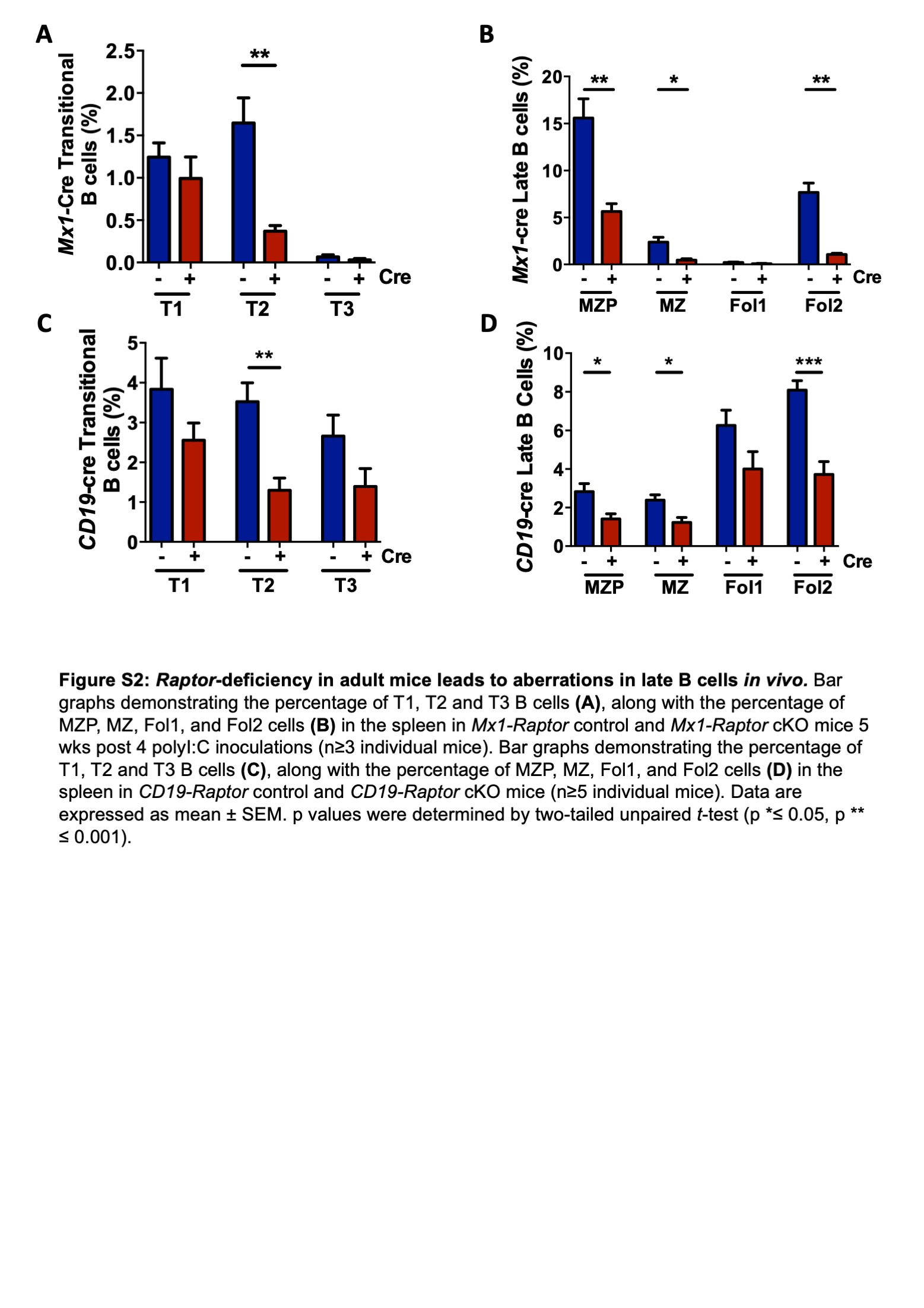

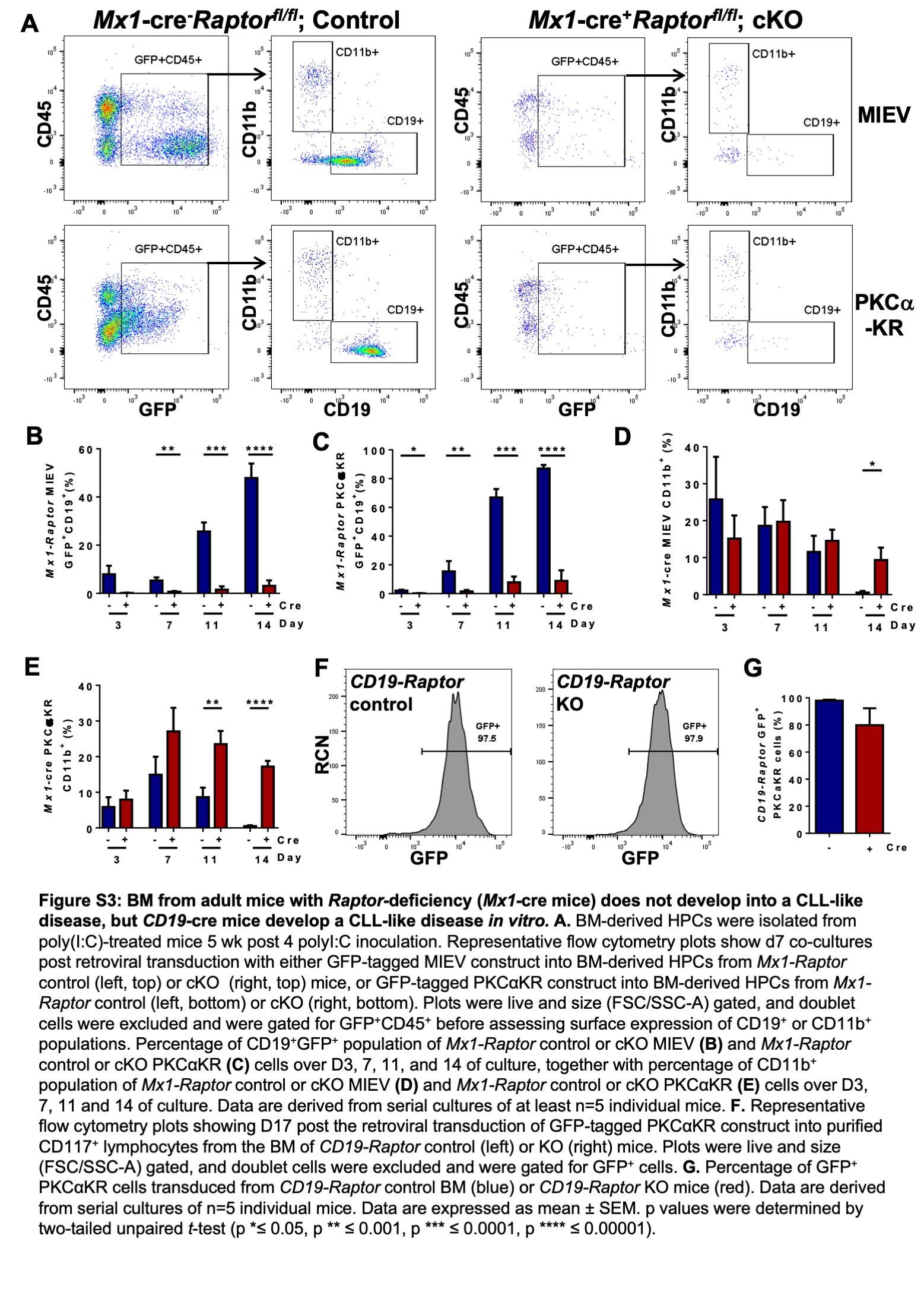

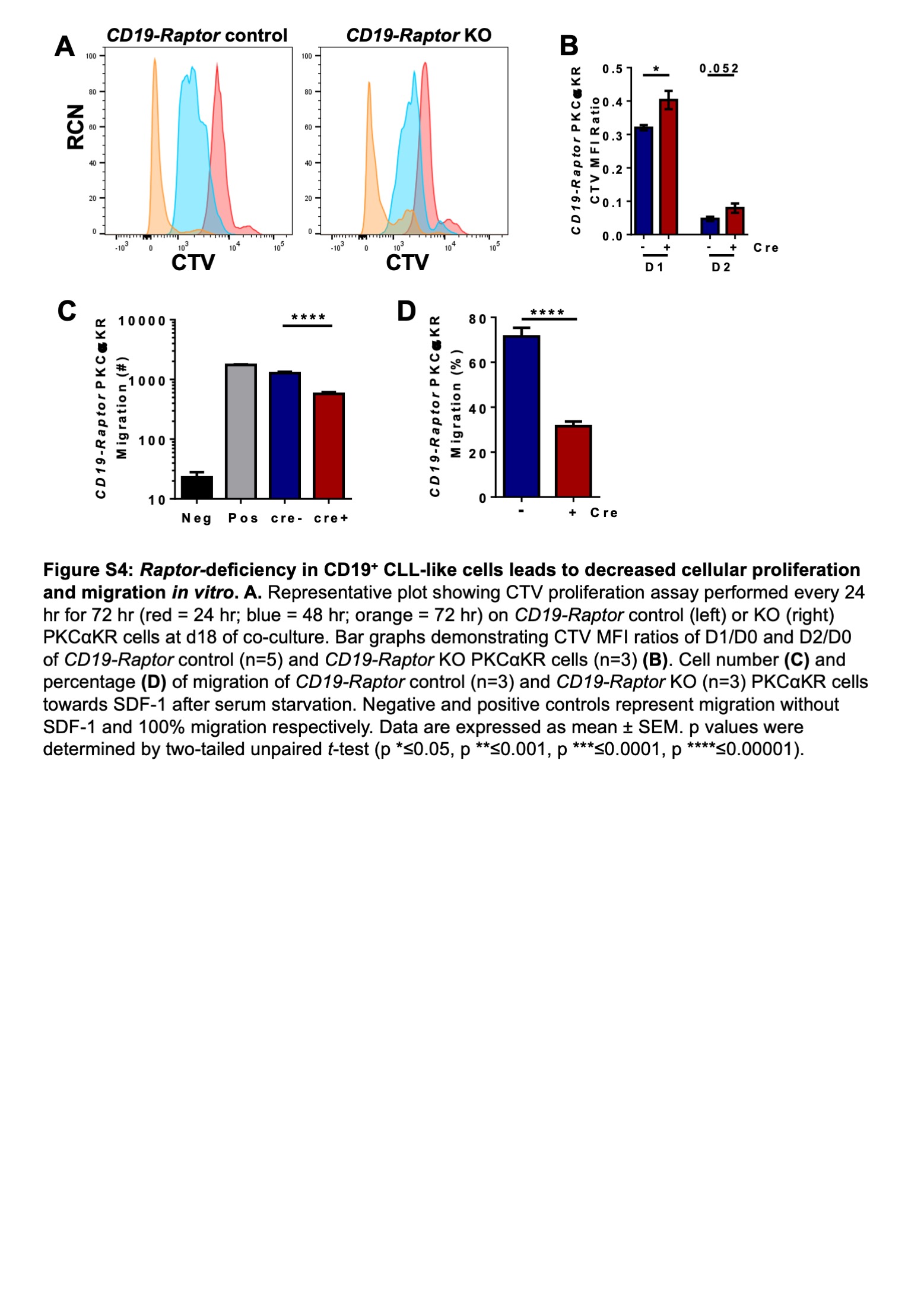

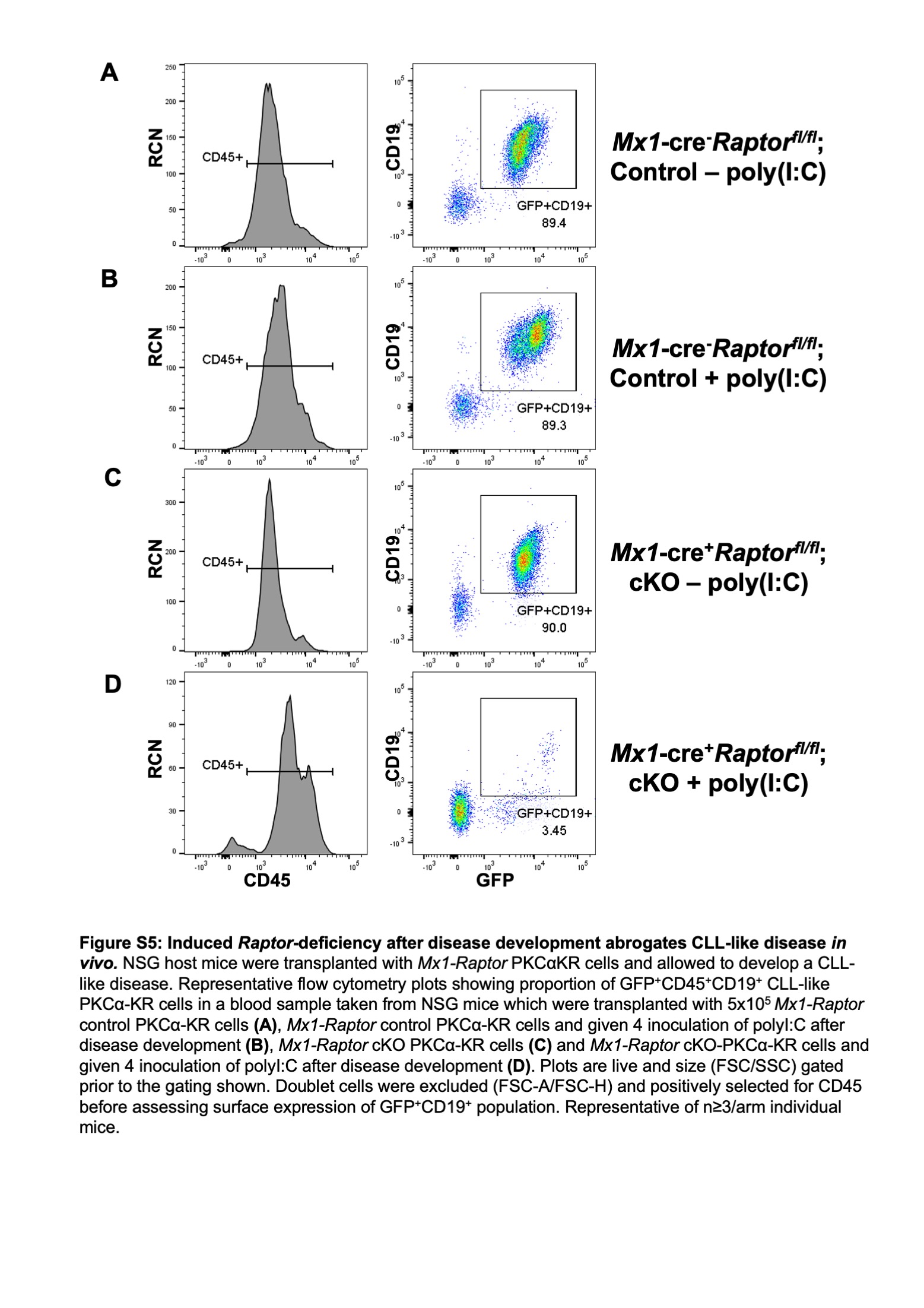

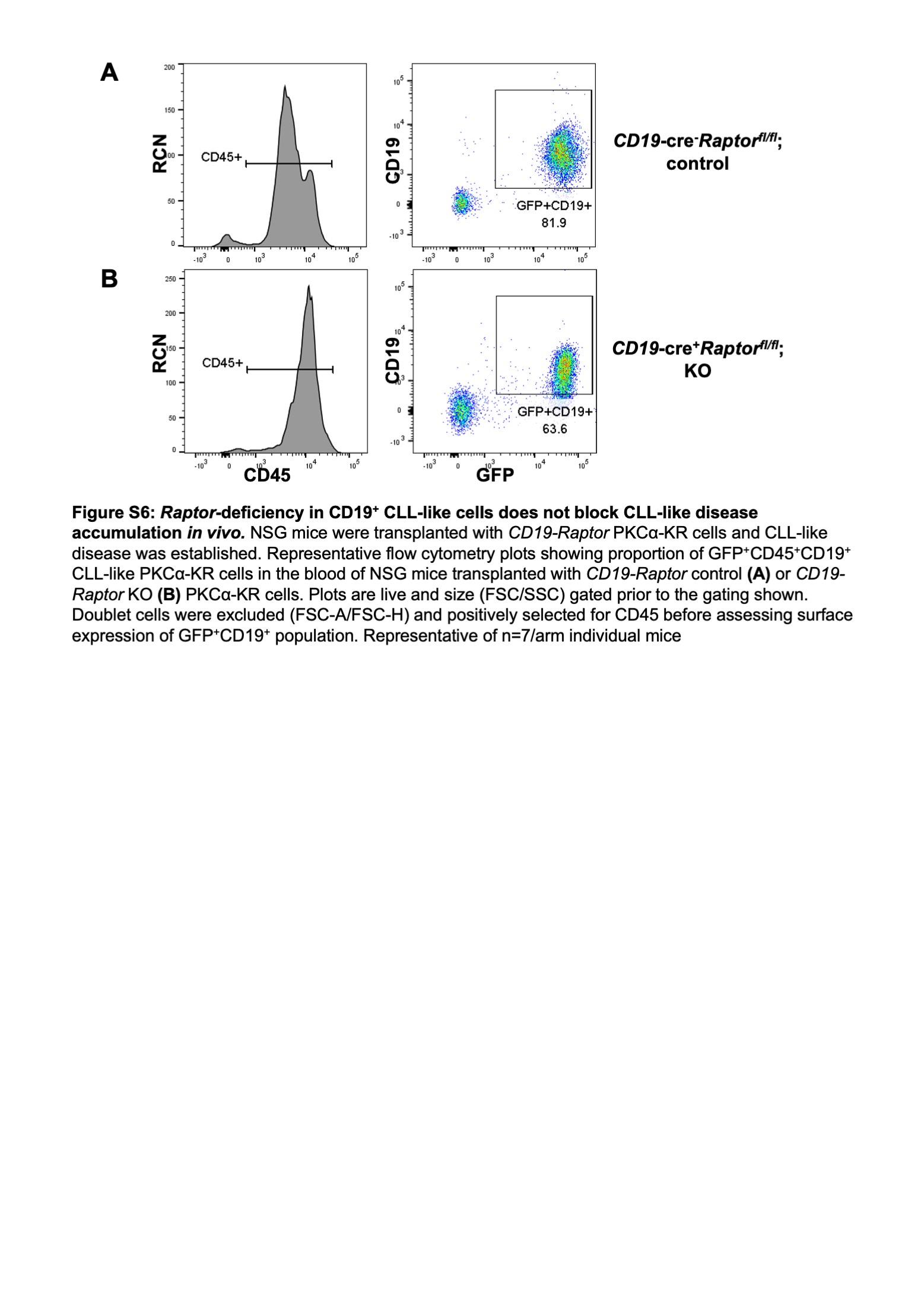

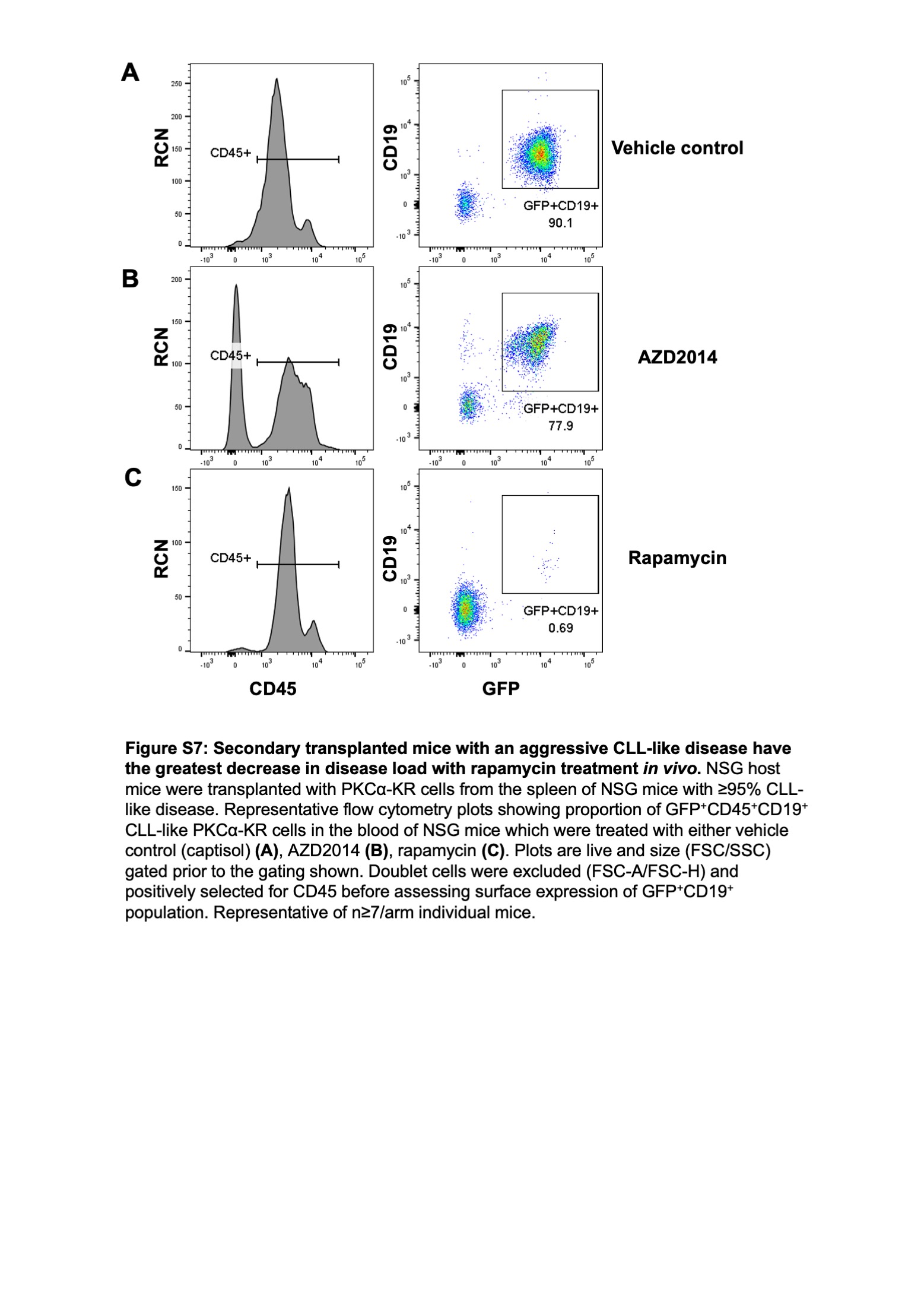

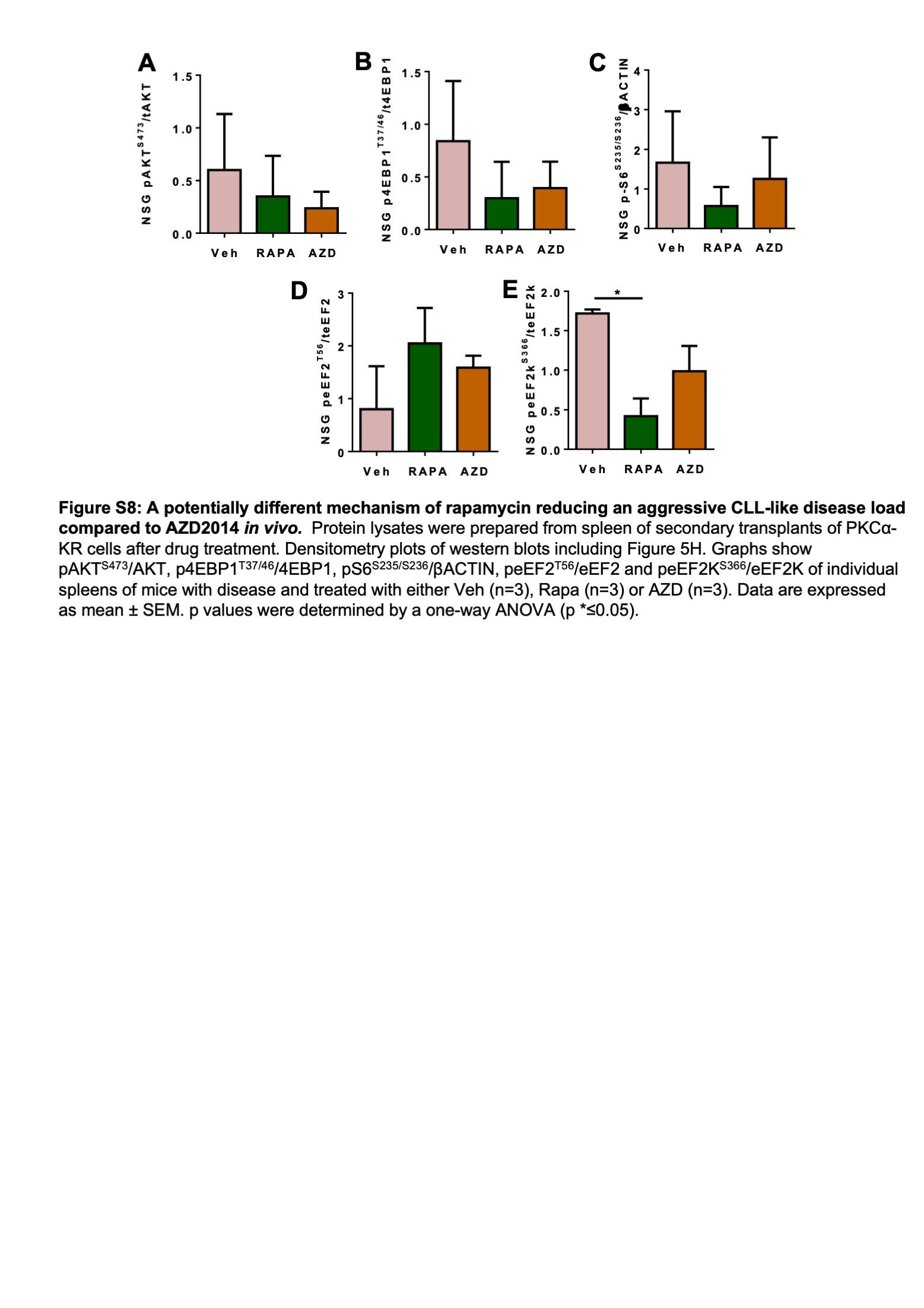

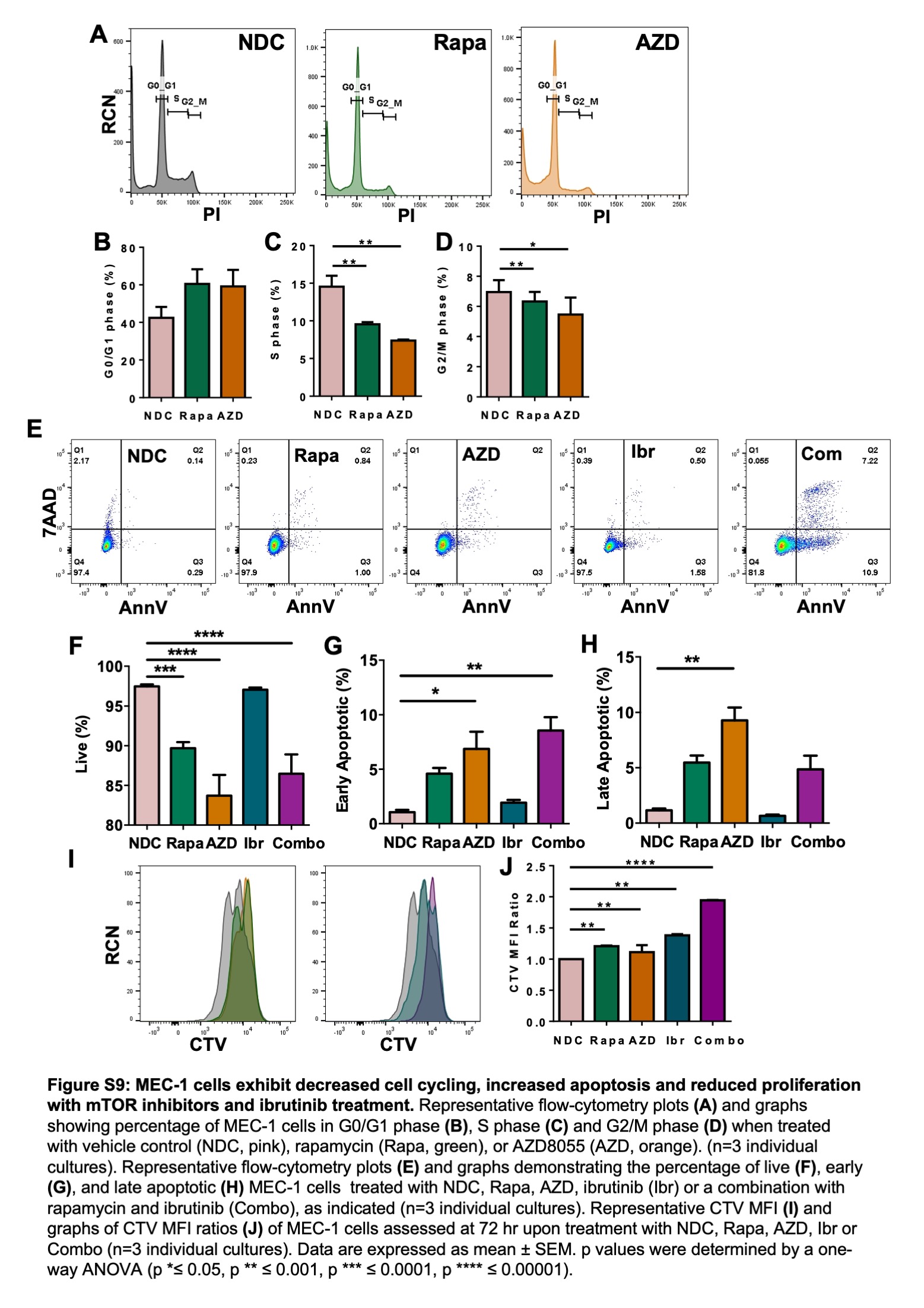

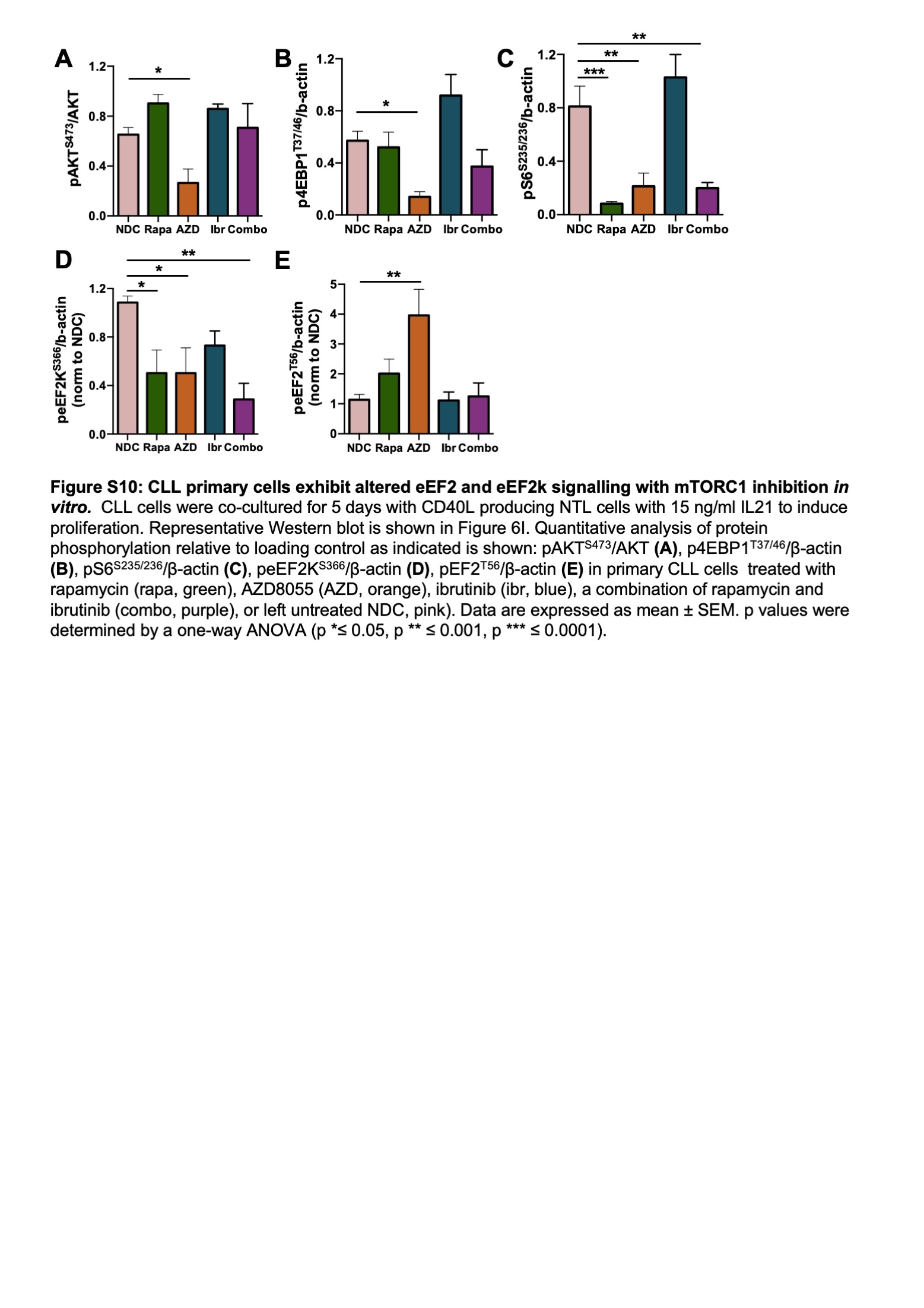

| **CLL ID** | **Treatment^a^** | **Sex** | **Binet Stage** | **ZAP-70 status^b^** | **Cytogenetics** |
| --- | --- | --- | --- | --- | --- |
| 113 | Yes | F | C | high | del17p |
| 140 | No | F | A | low | T12 |
| 147 | No | M | C | high | No 11q/17p |
| 151 | No | M | B | ND | del11q |
| 165 | No | M | C | ND | del11q |
| 177 | No | M |  | ND | No 11q/17p |
| 179 | Yes | M | B | ND | No 11q/17p |

Table S1. CLL patient clinical characteristics.

^a^ If previously undergone treatment, it was not within three months of sample collection.
^b^ ZAP-70 analysis was conducted by immunohistochemistry in the regional haematology laboratory. ND – not determined.

| Gene | Forward | Reverse | Species | Sequence Information |
| --- | --- | --- | --- | --- |
| *Ebf1* | tacagaaggtcattcctcgg | atcccatacagggcttcaac | Mouse | NM_001290709.1 |
| *Pax5* | acagga catggaggag tgaa | tgacaccttg atgggcaagt | Mouse | NM_008782.2 |
| *Rptor* | atggtagcaggcacactcttcatg | gctaaacattcagtccctaatc | Mouse | Ref. 50 |
| *Gusb* | taagacgctgatcacccaca | cagataacatccacgtacgg | Mouse | NM_010368.1 |
| *Tbp* | gtacccttcaccaatgactc | cagccaagattcacggtaga | Mouse | NM_013684.3 |

Table S2: List of primers used for PCR reactions.

The full sequence for each gene was obtained from the NCBI gene database (https://ncbi.nlm.nih.gov/gene). Each primer was designed to have close to 10 C=G and 10 A=T bonds. The length between the forward and reverse primer is between 150-300 base pairs. *Gusb* and *Tbp* were used as reference genes for murine cells. The specificity of each primer sequence was checked by using BLASTn tool.

| Name | Reactive Species | Clone | Dilution | 2^nd^ary Ab |
| --- | --- | --- | --- | --- |
| RAPTOR | Human, Mouse, Rat | 24C12 | 1:1000 | Rabbit |
| pAKT^S473^ | Human, Mouse, Rat | 23C8D2 | 1:1000 | Rabbit |
| AKT (pan) | Human, Mouse, Rat | C67E7 | 1:1000 | Rabbit |
| pS6^S235/S236^ | Human, Mouse, Rat | D57.2.2E | 1:1000 | Rabbit |
| S6 | Human, Mouse, Rat | 54D2 | 1:1000 | Mouse |
| p4EBP1^T37/T46^ | Human, Mouse, Rat | 236B4 | 1:1000 | Rabbit |
| pRb^S807/811^ | Human, Mouse, Rat |  | 1:1000 | Rabbit |
| Rb | Human | 4H1 | 1:1000 | Mouse |
| peEF2^T56^ | Human, Mouse, Rat |  | 1:1000 | Rabbit |
| eEF2 | Human, Mouse, Rat |  | 1:1000 | Rabbit |
| peEF2k^S366^ | Human, Mouse, Rat |  | 1:1000 | Rabbit |
| eEF2k | Human, Mouse, Rat |  | 1:1000 | Rabbit |
| Cyclin A |  |  | 1:1000 | Rabbit |
| 4EBP1 | Human, Mouse, Rat | 53H11 | 1:1000 | Rabbit |
| GAPDH | Human, Mouse, Rat | D16H11 | 1:1000 | Rabbit |
| β-ACTIN | Human, Mouse, Rat | C4 | 1:1000 | Mouse |
| α-mouse IgG, HRP Ab |  |  | 1:10000 |  |
| α-rabbit IgG, HRP Ab |  |  | 1:10000 |  |

Table S3: List of antibodies used for western blotting.

List of antibodies (Ab) and their dilutions in 5% BSA in TBS-T. All antibodies were purchased from Cell signalling (Herts, UK), except Cyclin A (Santa Cruz, USA) and β-ACTIN (Santa Cruz, USA).
